# Atlas of *DES* (desmin) variants: Impact of variants located within the head domain on filament assembly

**DOI:** 10.1101/2023.08.11.552974

**Authors:** Sabrina Voß, Volker Walhorn, Stephanie Holler, Anna Gärtner, Greta Pohl, Jan Gummert, Dario Anselmetti, Hendrik Milting, Andreas Brodehl

## Abstract

Desmin is a muscle-specific intermediate filament protein, which plays a significant role in providing structural integrity of cardiomyocytes by connecting different cell organelles and multi-protein complexes. *DES* mutations cause cardiomyopathies and skeletal myopathies. Most of these pathogenic mutations are localized in the highly conserved rod domain and affect the filament assembly.

However, the impact of *DES* variants within the N-terminal head domain on the filament assembly process is widely unknown. Therefore, we inserted a set of 85 different head domain variants with unknown significance from human genetic databases in expression constructs and investigated their impact on filament formation in cell culture in combination with confocal microscopy. The majority of these desmin variants do not affect the filament assembly. However, the desmin variants -p.S13P, -p.N107D, -p.E108G and -p.K109E significantly inhibit the filament assembly. Additionally, we expressed and purified recombinant desmin and investigated the filament assembly defects by atomic force microscopy verifying these findings at the single molecular level. Furthermore, we truncated systematically the head domain to investigate which general parts of this domain are necessary for filament assembly.

In summary, our functional investigations might be relevant for the classification of novel *DES* variants and the genetic counselling of patients carrying desmin head variants.

## Introduction

Desminopathies (OMIM, #125660; ORPHA:98909) are caused by mutations in the *DES* gene, encoding the intermediate filament (IF) protein desmin. Desmin is the major muscle IF protein and is expressed in smooth muscle cells, skeletal myocytes and cardiomyocytes [1, 2]. IFs connect different multi-protein complexes like the cardiac desmosomes [3–5], costameres [6, 7], Z-bands [8] and different cell organelles like e.g. the mitochondria [9] and the nuclei [10]. Therefore, desmin is highly important for the structural integrity of (cardio)myocytes.

Clinically, desminopathies manifest as different cardiomyopathies and/or skeletal myopathies [11–14]. The clinical spectrum of desminopathies is broad and even within the same family different types of cardiomyopathies may result [15]. Ventricular and atrial arrhythmias are frequently observed in patients with desminopathies [16–19] leading in some cases to cardiac arrest or even sudden cardiac death [20, 21].

Most of the pathogenic *DES* mutations are heterozygous missense or small in-frame deletion mutations [22] causing toxic desmin aggregates [23–25]. Interestingly, expression of mutant desmin causes a co-aggregation of the wild-type form indicating a dominant negative effect on filament assembly [26–28]. In contrast, C-terminally truncating desmin mutations are rare and are only pathogenic in homozygous or compound heterozygous genotypes [29–32].

Desmin consists like all IF proteins of a central α-helical rod domain flanked by non-helical head and tail domains [33]. The rod domain is subdivided into the 1A, 1B and 2 coil subdomains connected by non-helical linkers [34]. Two monomers dimerize by formation of coiled-coil dimers [35]. These dimers anneal into anti-parallel tetramers [36] and unit-length filaments (ULFs) are formed by lateral association of several tetramers [37]. The ULFs are the essential building blocks of the IFs [38]. In addition, IFs can fuse at their ends [39] and exchange subunits alongside their longitudinal direction indicating a highly dynamic process [40].

Of note, desmin mutations interfere at different stages within the complex filament formation [23]. We have recently identified a genetic hotspot in the N-terminal part of the 1A subdomain where several likely pathogenic mutations affect the desmin filament assembly [41]. Up to now, some desmin head mutations have been previously described [15, 42–44]. However, most of these desmin variants are classified as variants of unknown significance (VUS) in genetic databases.

Here, we generated a set of 85 different rare missense VUS localized in the head domain of desmin and analyzed their effects on filament formation in cell culture using confocal microscopy. Our experiments indicate that the majority of mutations in the head domain do not interfere with desmin filament assembly. However, the variants *DES*-p.S13P, -p.N107D, - p.E108G and -p.K109E cause a cytoplasmic desmin aggregation. Of note, p.N107D, p.E108G and p.K109E are localized in close proximity to the 1A domain, which was recently recognized as a mutation hotspot for pathogenic desmin mutations [41]. These data were verified at the single molecular level by atomic force microscopy (AFM). In addition, we generated a set of different N-terminal truncating mutations to investigate the general relevance of the head domain for desmin filament assembly. These experiments demonstrate that larger N-terminal deletions cause a filament assembly defect.

In conclusion, our *in vitro* experiments systematically provide functional insights about desmin head domain variants and can contribute in future to variant classification. Therefore, these functional data might be relevant for the genetic diagnostics of patients carrying *DES* mutations within the head domain.

## Material and Methods

### Genetic Disease Database Analysis

85 rare missense VUS affecting amino acids within the desmin head domain were selected from the Human Gene Mutation Database (https://www.hgmd.cf.ac.uk/) and the ClinVar Database (https://www.ncbi.nlm.nih.gov/clinvar/, accessed on 10^th^ January 2023). A minor allele frequency (MAF) <0.0001 (Genome Aggregation Database, https://gnomad.broadinstitute.org/) was used as an exclusion criterion since it is above the prevalence of cardiomyopathies.

### Cloning and Site-Directed Mutagenesis

The plasmid pEYFP-N1-DES-WT was generated by ligation of the human *DES* cDNA via *Xho*I and *BamH*I restriction sites as previously described [27] (Supplements, Figure S1A). Cloning of pmRuby-N1-DES-WT was previously described [26] (Supplements, Figure S1B). The bacterial expression plasmid pET100D-TOPO-DES was generated by TOPO cloning according to the manufacturer’s instructions (Thermo Fisher Scientific, Waltham, MA, USA) and was previously described [27] (Supplements, Figure S1C). Most missense and deletion mutations were inserted by site-directed mutagenesis using the QuikChange Lightning Kit (Agilent Technologies, Santa Clara, CA, USA) in combination with appropriate oligonucleotides (Supplements, Table S1, synthesized by Microsynth, Balgach, Switzerland) according to the manufacturer’s instructions. *Escherichia coli* (*E. coli*, XL-10 gold) bacteria were transformed by heat shock (30 sec, 42 °C) and colonies were selected overnight using 50 µg/mL kanamycin sulfate (Sigma-Aldrich, Darmstadt, Germany). Some N-terminal deletion mutations were cloned by PCR using Phusion Polymerase (Thermo Fisher Scientific) followed by ligation using T4 DNA ligase (Thermo Fisher Scientific) (Supplements, Table S1). Plasmids were prepared from single colonies using the GeneJET Plasmid Miniprep Kit (Thermo Fisher Scientific) as described by the manufacturer. All plasmids were verified in the desmin coding region by Sanger sequencing (Macrogen, Amsterdam, Netherlands) using the CMV-for and EGFP-rev primers for the pEYFP-N1-DES constructs or the T7-for and T7-rev primers for the pET100D-TOPO-DES constructs (Supplements, Table S1). The sequencing data were analyzed using SnapGene 6.1 software (GSL Biotech, San Diego, CA, USA).

### Cell Culture

Dulbecco’s Modified Eagle Medium (DMEM, Thermo Fisher Scientific) was used for culturing of H9c2 and SW-13 cells (ATCC, Manassas, VA, USA) at 37 °C and 5 % CO_2_. DMEM was supplemented with 10 % fetal calf serum and penicillin/streptomycin. Cells were subcultured at a confluency of >90 % using trypsin/ethylendiaminetetraacetic acid (Thermo Fisher Scientific).

Human induced pluripotent stem cells (iPSCs) generated from a healthy donor (NP0040-8, UKKi011-A, kindly provided by Dr. Tomo Saric, University of Cologne, Germany) were cultured in Essential 8 Medium (Thermo Fisher Scientific) on vitronectin-coated plates. Versene solution (Thermo Fisher Scientific) was used for cell dissociation.

### Cardiomyocyte Differentiation of Induced Pluripotent Stem Cells

iPSCs were differentiated into cardiomyocytes by modulation of the *Wnt*-pathway as previously described [45]. Contracting iPSC-derived cardiomyocyte clusters (Supplements Video File S1) were imaged with the Eclipse TE2000-U wide-field microscope in combination with the Plan Fluor 4x/0.13 PhL DL objective and the Lucia software (Nikon, Tokyo, Japan).

### Cell Transfection

24 h before transient transfection, SW-13 and H9c2 cells were subcultured and 200.000 cells were cultured in 8-well µSlide chambers (ibidi, Gräfelfing, Germany) or in 6-well plates for expression analysis. Beating iPSC-derived cardiomyocytes were split using Accutase / trypsin (1:1; Sigma-Aldrich) for 6 min at 37 °C, centrifuged and cultured on vitronectin-coated (Sigma-Aldrich) µSlide chambers. The cells were transiently transfected using Lipofectamin 3000 (Thermo Fisher Scientific) according to the manufacturer’s instructions. Afterwards, the cells were cultured for 24 h at 37 °C and 5 % CO_2_.

### Cell Fixation and Immunocytochemistry

Cells were washed with phosphate buffered saline (PBS, Thermo Fisher Scientific). 4 % HistoFix (Carl Roth, Karlsruhe, Germany) was used for cell fixation (15 min at room temperature, RT). After two washing steps with PBS, the cells were permeabilized using 0.1 % Triton X-100 (15 min, RT). Subsequently, the cells were washed with PBS. Phalloidin conjugated with the fluorescence dye Texas Red (1:400, Thermo Fisher Scientific) was used for staining of F-actin (40 min, RT), 4′,6-diamidino-2-phenylindole (DAPI, 1 µg/mL) was used for staining of the nuclei (5 min, RT). α-Actinin was stained by using antibodies as recently described [41]. Finally, two washing steps with PBS were performed and the cells were stored at 4 °C until confocal microscopy analyses.

### Confocal Microscopy

Confocal laser scanning microscopy was performed using the TCS SP8 system (Leica Microsystems, Wetzlar, Germany) as previously described [46]. Briefly, DAPI, EYFP and Texas Red were excited at 405, 488 and 552 nm. The fluorescence emissions of these dyes were sequentially detected between 410-460, 493-560 and 570-758 nm. The fluorescent protein mRuby [47] was excited at 552 nm and the fluorescence emission was detected between 557-643 nm. 3D stacks were recorded and processed using the Las X software (Leica Microsystems). All representative cell images are shown as maximum intensity projections.

### Co-localization Analysis

Co-localization of desmin-EYFP and desmin-mRuby constructs was evaluated using the Fiji software in combination with the EzColocalization plugin [48]. The Pearson correlation coefficients (PCC) of >10 double transfected cells were determined.

### Expression Analysis

Expression analysis of desmin-EYFP was done by fluorescence intensity measurements using the Infinite M1000 plate reader (Tecan, Männedorf, Switzerland) at 37 °C. Transfected cells were washed twice with PBS and were afterwards excited at 488 nm. Fluorescence emission was determined at 505 nm with a bandwidth of 20 nm. Four independent transfection experiments per construct were analyzed and normalized by subtraction of background signals of non-transfected cells (NT).

### Recombinant Desmin Expression

*E. coli* (BL21 Star DE3, Thermo Fisher Scientific) were transformed with wild-type or mutant pET100D-TOPO-DES expression plasmids by heat shock treatment (30 sec, 42 °C). After selection using 100 µg/mL ampicillin over night, single colonies were inoculated and expanded in Lysogeny Broth medium (Carl Roth). Desmin expression was induced by isopropyl β-D-thiogalactoside (1 mM) for 4 h under vigorous shaking at 37 °C. Afterwards, bacteria were collected by centrifugation and stored at -80 °C.

### Purification of Recombinant Desmin

Bacteria were lysed and the inclusion bodies were prepared as previously described [27]. Afterwards, recombinant desmin was purified under denaturing conditions (8 M urea) by ion exchange chromatography (IEC) using 5 mL HiTrap DEAE-FF columns (GE Healthcare, Chicago, IL, USA) and in a second step by immobilized metal affinity chromatography (IMAC) using HisTrap FF Crude columns (Cytiva, Marlborough, MA, USA) in combination with the ÄKTApurifier system (GE Healthcare). The purified recombinant desmin was stored at -80 °C.

### Atomic Force Microscopy

Recombinant desmin was dialyzed stepwise in buffer without urea (5 mM Tris-HCl, 1 mM dithiothreitol, pH 8.4) and diluted to a concentration of 0.3 g/L. The filament assembly was initiated by adding an equal volume of sodium chloride buffer (200 mM NaCl, 45 mM Tris-HCl, pH 7.0) and heating to 37 °C for 1 h [49]. Poly-L-ornithine solution (molecular weight 30.000-70.000, 0.01%, Sigma-Aldrich) was applied to freshly cleaved mica substrates (Plano, Wetzlar, Germany) and incubated for 5 min. Subsequently, the mica substrates were rinsed with deionized water to remove excess polypeptides. Readily assembled desmin was applied to the modified mica substrates and incubated for approximately 30 s. Unbound desmin was removed by rinsing with deionized water. AFM imaging was performed in tapping mode at RT in water using a JPK NanoWizard ULTRA Speed 2 (JPK Bruker, Berlin, Germany) and USC-F0.3-k0.3 Cantilevers (Nano World, Neuchâtel, Switzerland).

### Molecular Modelling and Sequence Alignment

AlphaFold Multimer was used for prediction of the human dimeric desmin structure [50]. The molecular structure was visualized by PyMOL Molecular Graphics Version 2.52. (Schrödinger, New York, NY, USA). The multiple sequence alignment was performed using Clustal Omega (https://www.ebi.ac.uk/Tools/msa/clustalo/, accessed on 11^th^ January 2023) [51] using the following desmin reference sequences: *Homo sapiens* (NP_001918.3), *Xenopus laevis* (NP_001080177.1), *Mus musculus* (NP_034173.1), *Rattus norvegicus* (NP_071976.2), *Danio rerio* (NP_571038.2 and NP_001070920.1).

### Statistical Analysis

In minimum, four independent transfection experiments were performed per construct and per cell line. In each transfection experiment, about 100 transiently transfected cells were analyzed. The percentage of aggregate formation was calculated for each independent experiment. The non-parametric Kruskal-Wallis test followed by Dunn’s multiple comparison was used for statistical analysis using GraphPad Prism Version 9.0 (GraphPad Software, San Diego, CA, USA). All data are shown as means ± standard deviations (SD).

## Results

Several amino acids within the head domain of desmin are highly conserved (Figure 1A) indicating their functional relevance. We analyzed the genetic disease databases ‘*ClinVar*’ (https://www.ncbi.nlm.nih.gov/clinvar/) and the ‘*Human Gene Mutation Database’* (https://www.hgmd.cf.ac.uk/) on disease related variants. Desmin VUS localized in the head domain were included if their MAF were below 0.0001 (according to the *Genome Aggregation Database*, https://gnomad.broadinstitute.org/). A set of 85 different mutant expression constructs was generated (Figure 1B, Figure S1A). Afterwards, we transfected SW-13 cells with these different desmin expression constructs and analyzed the filament formation by confocal microscopy. We used SW-13 cells, since they do not express endogenous desmin or any other cytoplasmic IF proteins [52]. Wild-type desmin-EYFP formed filamentous structures of different size and shape (Figure 1C) comparable as previously described [53]. Previously, we have shown that the C-terminal fluorescent protein tag does not affect the filament assembly of desmin [27]. The majority of desmin VUS in the head domain did not interfere with the filament assembly since they form in most cases filamentous structures comparable to the wild-type desmin (Figure 2). However, we identified four *DES* variants affecting the filament assembly process. Desmin-p.S13P and -p.N107D formed in the majority of transfected cells (80.7 and 65.2 %) a mixed cellular phenotype consisting of filamentous structures and cytoplasmic aggregates within the same cell (Figure 2L, 2*d, Figure 3). For desmin-p.E108G and -p.K109E we detected cytoplasmic aggregate formation in all transfected SW-13 cells (Figure 2*e-*f). Statistical analysis verified these findings (Figure S2). Analysis of fluorescence intensities revealed no significantly different expression of these mutants in comparison to the wild-type desmin (Supplements, Figure S3).

**Figure 1.**
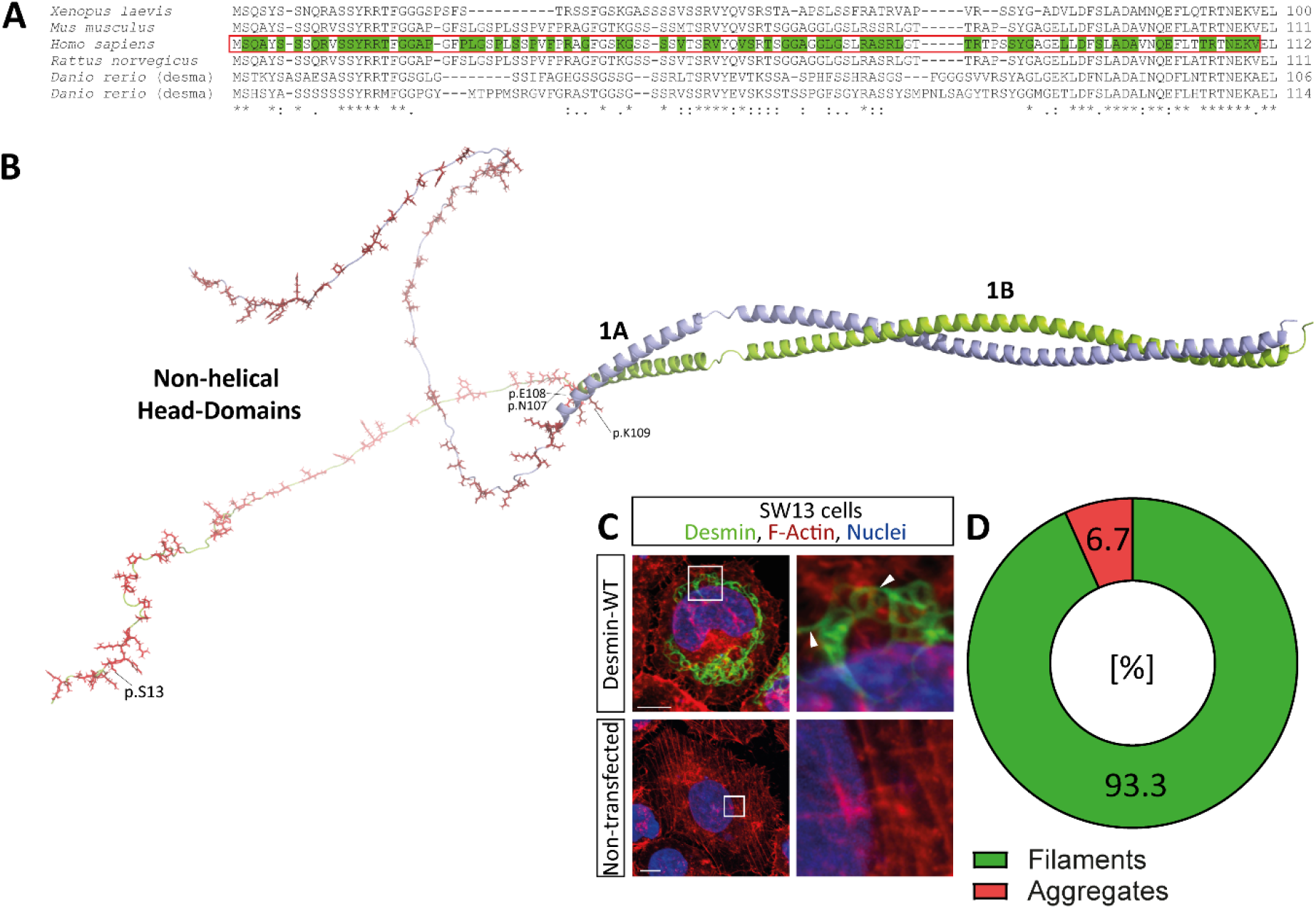
**(A)** Partial sequence alignment of the desmin head domains (red box) from different species (*Xenopus laevis*, *Mus musculus*, *Homo sapiens*, *Rattus norvegicus* and *Danio rerio*). Positions of variants with unknown significance (VUS) are highlighted in green. Fully conserved amino acids are indicated with stars and conserved substitutions with colons. Dots mark conserved positions with weakly similar properties of these amino acids. **(B)** Structural overview of the non-helical head and the α-helical 1A and 1B domains of the desmin dimer modelled using AlphaFold. Amino acids affected by VUS are shown as red sticks. **(C)** Representative maximal intensity projections of SW-13 cells expressing wild-type desmin (WT, green) and not-transfected controls. F-actin is shown in red and the nuclei are shown in blue. Scale bars represent 10 µm. Of note, desmin-WT forms filamentous structures. **(D)** Pie chart of observed cellular phenotypes (filament or aggregate formation) based on five independent transfection experiments of desmin-WT expressing SW-13 cells. Desmin-WT forms in the majority of transfected cells (93.3 %) filamentous structures.

**Figure 2.**
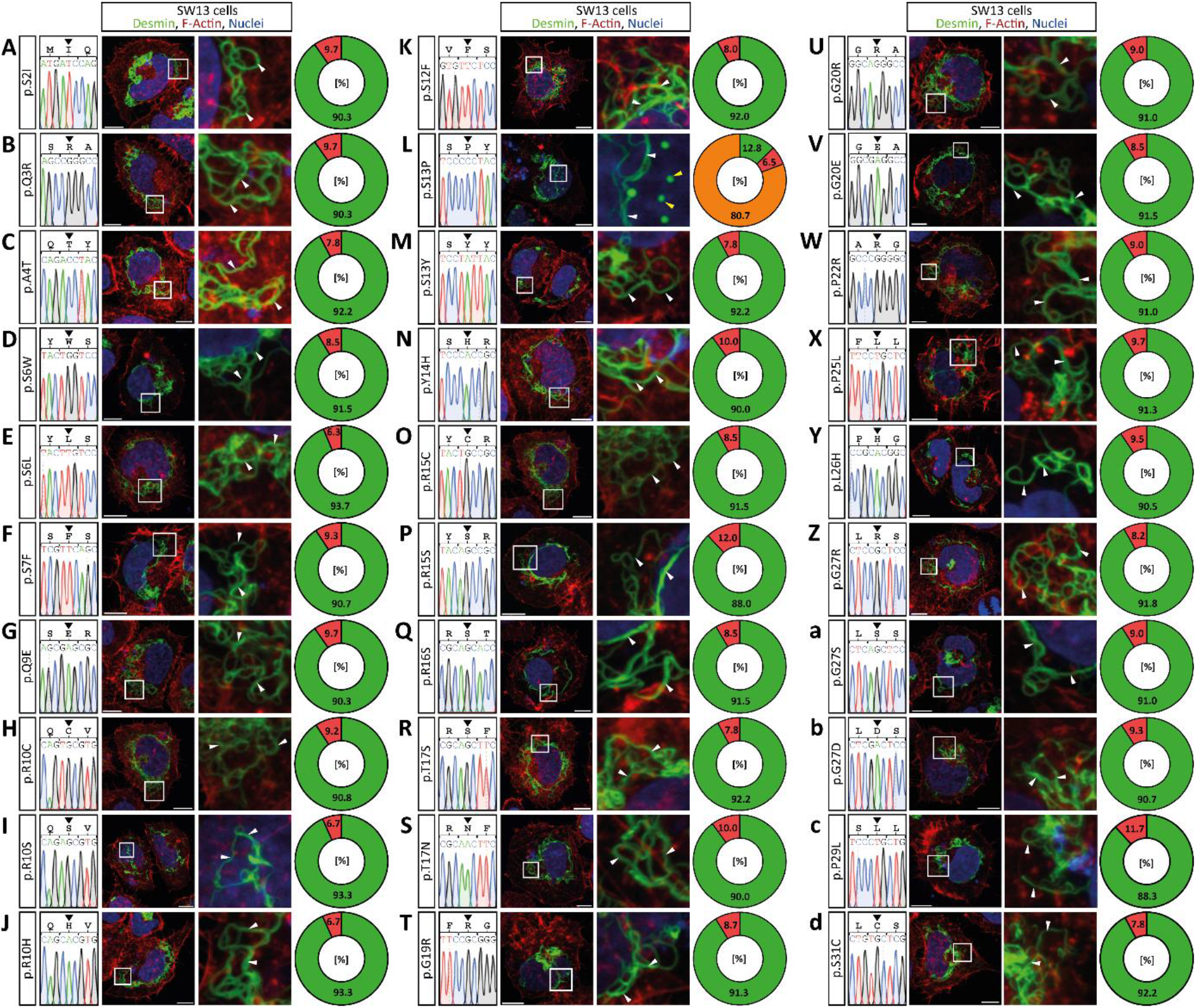
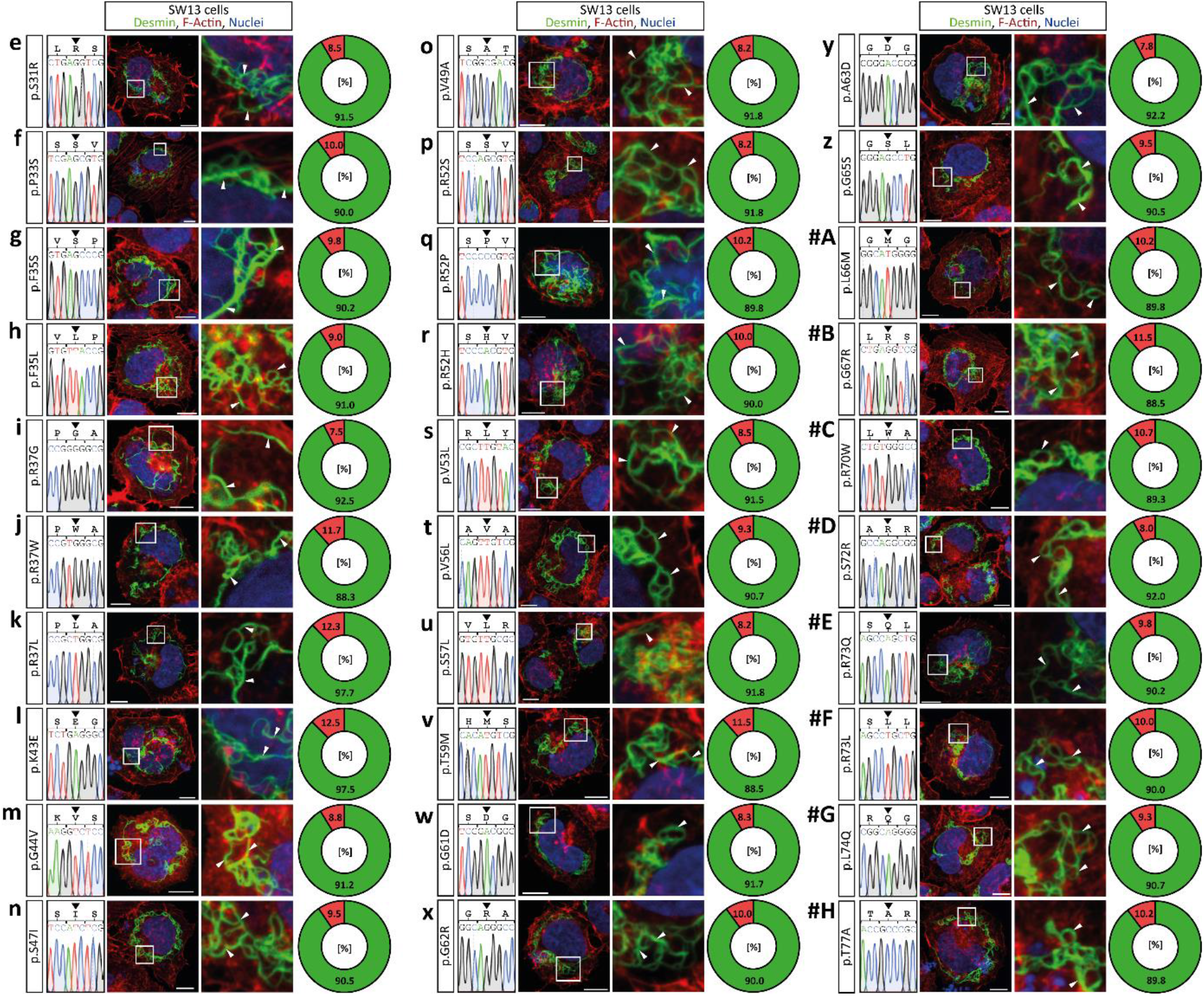
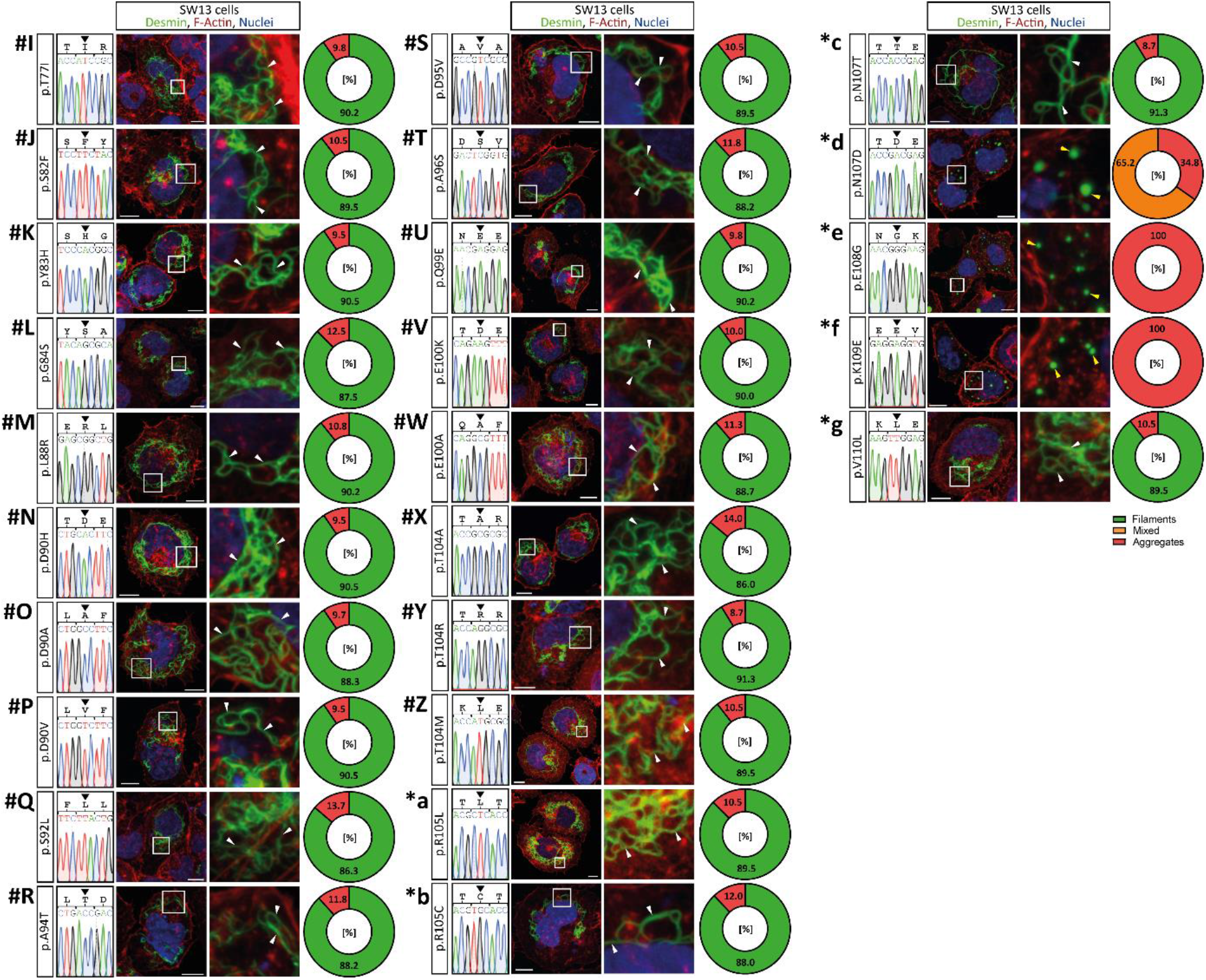
**(A-*g)** Partial electropherograms of the 85 generated VUS constructs and representative maximal intensity projections of transfected SW-13 cells expressing mutant desmin are shown (green). F-actin was stained using phalloidin-Texas Red (red) and the nuclei were stained using DAPI (blue). Scale bars represent 10 nm. The percentage of the different cell phenotypes is summarized as pie charts. The majority of desmin mutants form filamentous structures comparable to the wild-type desmin. Of note, two desmin mutants (p.S13P and p.N107D) form in most cells mixed phenotypes **(L and *d)** and two other desmin mutants (p.E108G and p.K109E) form predominant cytoplasmic aggregates **(*e and *f)**. The yellow arrow heads indicate cytoplasmic aggregates and the white arrow heads indicate filamentous structures.

**Figure 3.**
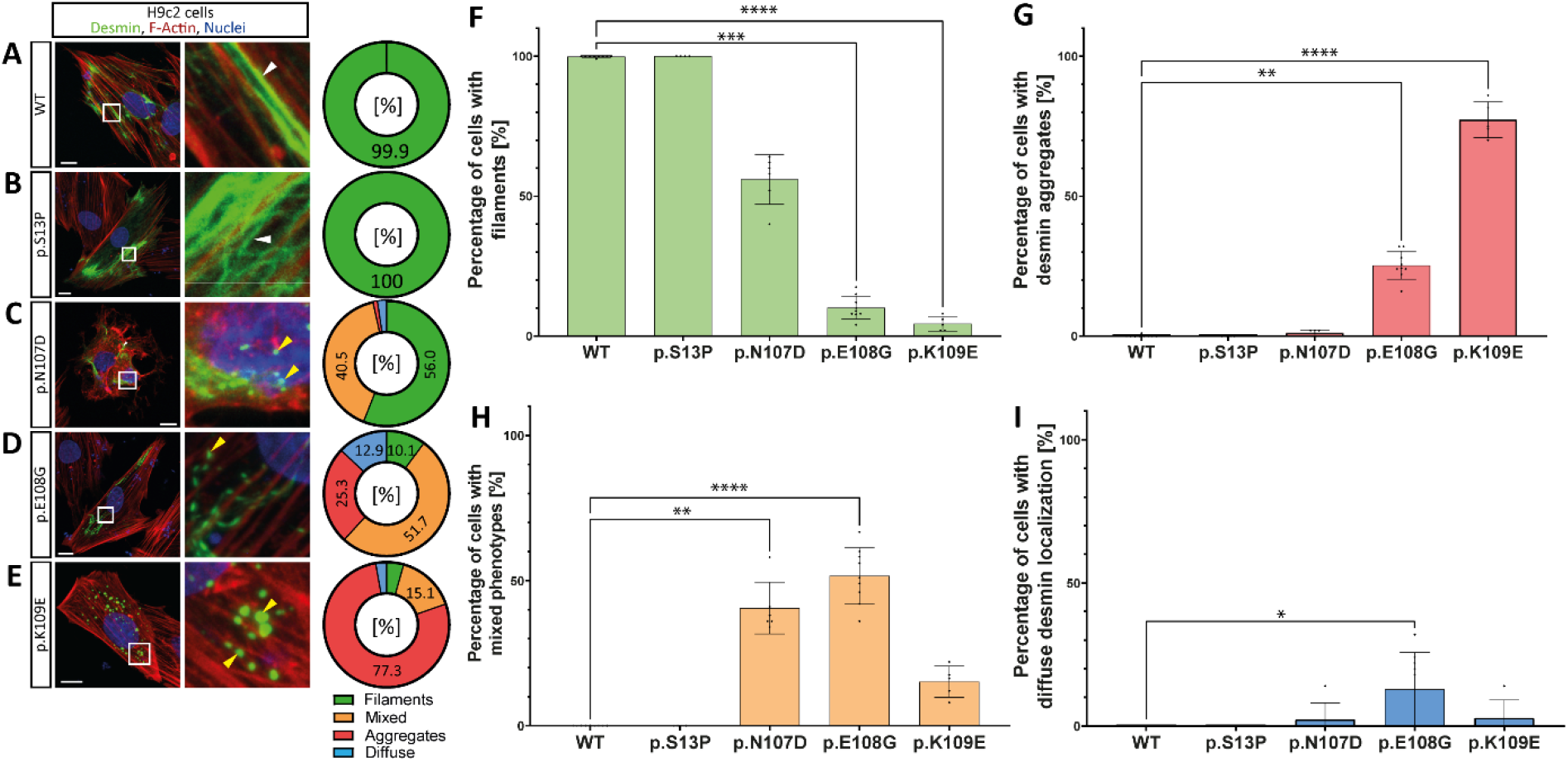
Representative images of H9c2 cells expressing wild-type or mutant desmin are shown **(A-E).** Desmin is shown in green, F-actin in red and the nuclei in blue. Scale bars represent 20 µm. Filaments are indicated with white and aggregates with yellow arrow heads. The percentages of the different cell phenotypes are summarized as pie charts. Statistical analysis was performed using non-parametric Kruskal-Wallis test **(F-I)**. All data are shown as mean ± SD.

To verify these experiments, we transfected H9c2 myoblasts and cardiomyocytes differentiated from iPSCs since both cell types express endogenous desmin. In case of desmin-p.N107D and -p.E108G the majority of H9c2 cells showed a mixed phenotype (Figure 3). Most of the desmin-p.K109E cells presented a desmin aggregation phenotype (Figure 3E-I). Of note, no desmin aggregation was detected for mutant p.S13P (Figure 3B) in H9c2 cells. These cells showed comparable filamentous desmin structures as in wild-type transfected cells (Figure 3A-B).

Comparable to this finding, desmin-p.E108G and -p.K109E formed cytoplasmic aggregates in 100 % of transfected iPSC-derived cardiomyocytes (Figure 4), whereas mutant desmin-p.S13P and -p.N107D formed in the majority of transfected cells filamentous structures comparable the wild-type control cells (Figure 4A-C).

**Figure 4.**
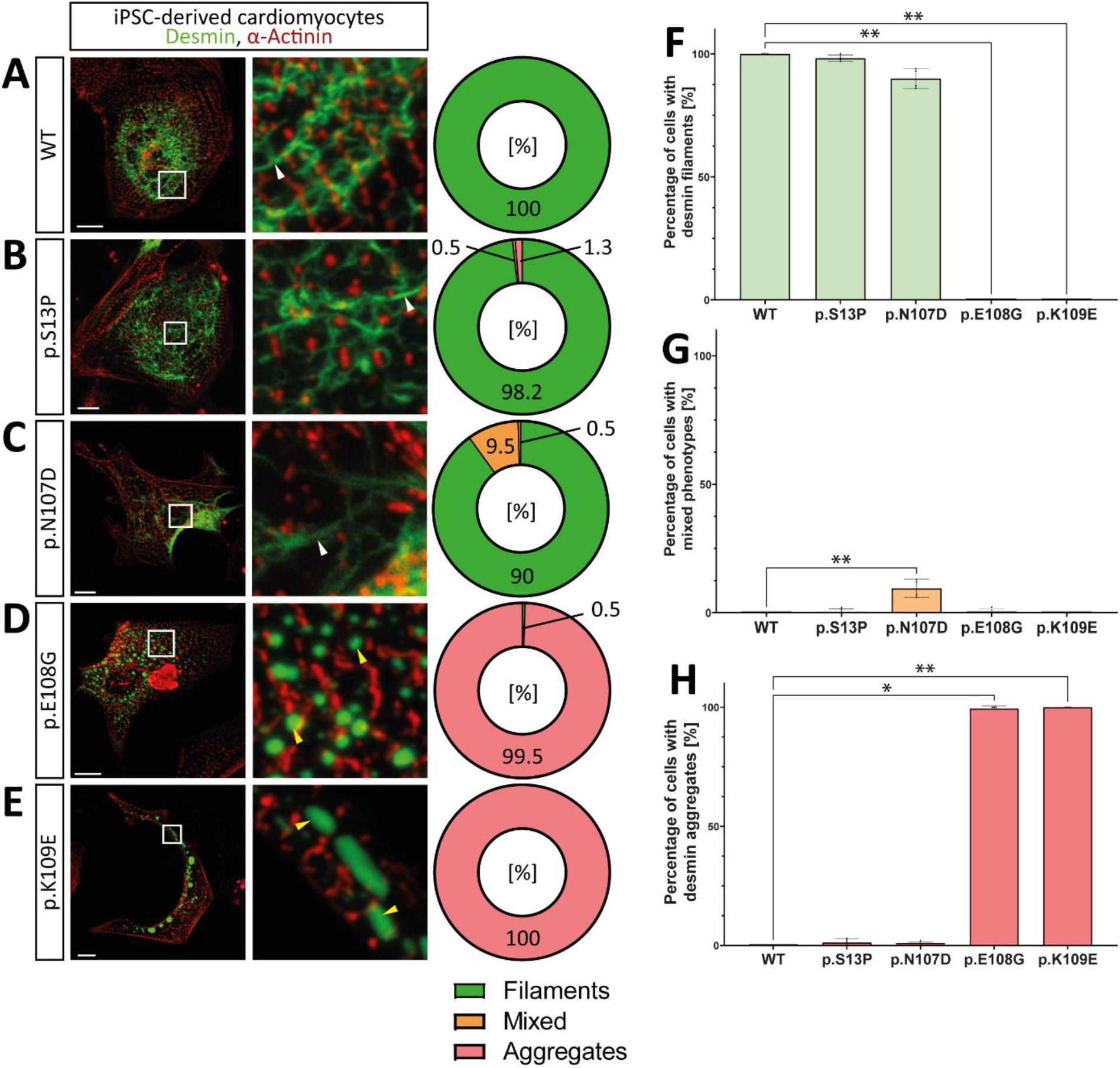
Representative images of iPSC-derived cardiomyocytes expressing wild-type or mutant desmin are shown **(A-E).** Desmin is shown in green, α-actinin, used as a cardiomyocyte marker, in red and the nuclei in blue. Scale bars represent 20 µm. Filaments are indicated with white and aggregates with yellow arrow heads. The percentages of the different cell phenotypes are summarized as pie charts. Statistical analysis was performed using non-parametric Kruskal-Wallis test **(F-H)**. All data are shown as mean ± SD.

Next, we expressed and purified wild-type and mutant recombinant desmin (Supplements, Figures S4 and S5) and analyzed *in vitro* the filament assembly by AFM (Figure 5A). Wild-type desmin formed filaments of different length (Figure 5B). In contrast, desmin-p.S13P formed only small structures, which are presumably ULFs based on their size (Figure 5C). The mutants p.N107D and p.K109E formed larger filamentous aggregates (Figure 5D and 5F). Desmin-p.E108G formed *in vitro* small fibrils and filaments with sharp kinks and edges (Figure 5E). These experiments revealed for all four desmin mutants abnormal desmin structures at the molecular level indicating an intrinsic filament assembly defect.

**Figure 5.**
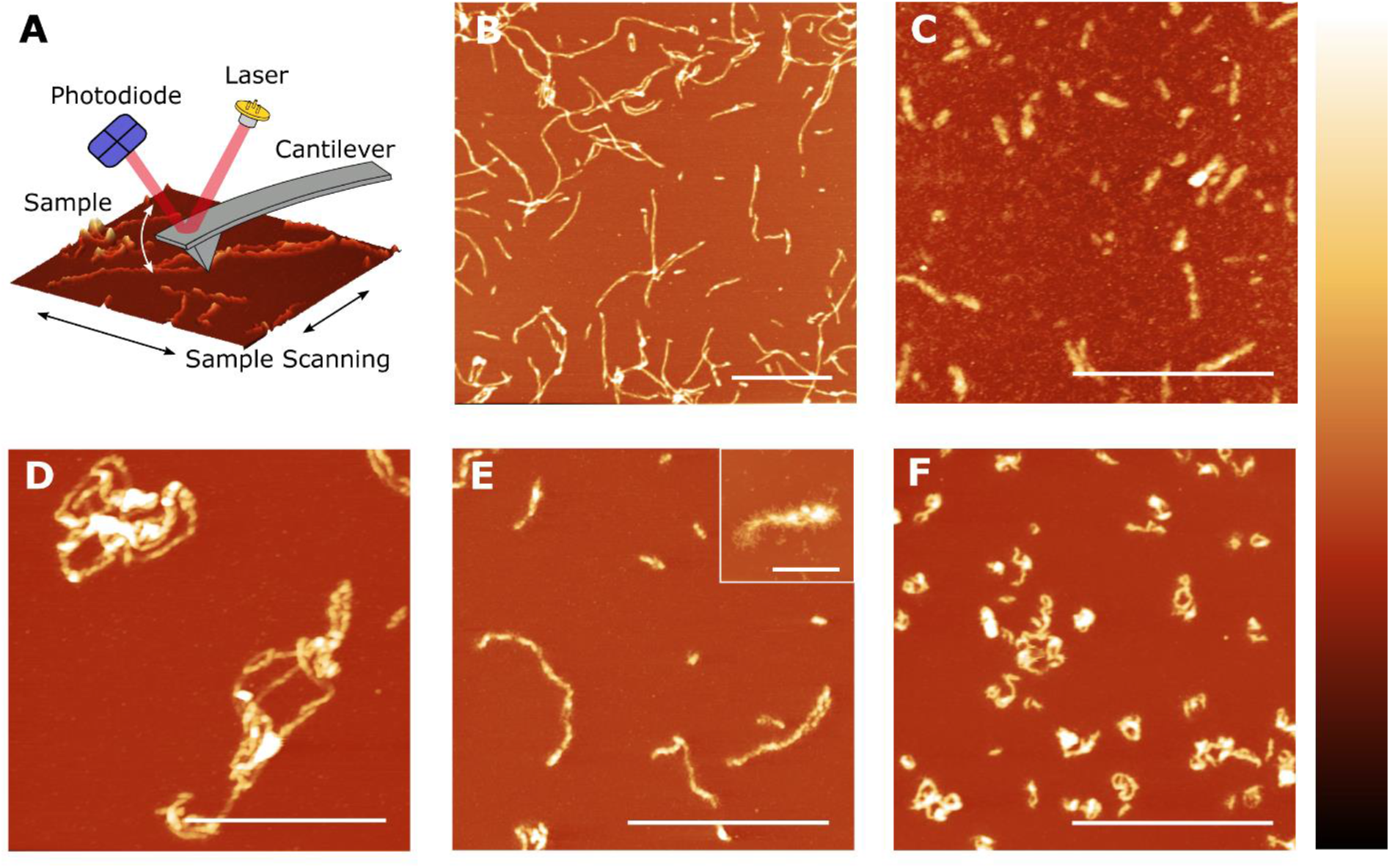
**(A)** Working principle of an AFM. (**B-F)** Typical topographical AFM scans of recombinant desmin after assembly. **(B)** Desmin wild-type exhibits long (several 100 nm) predominantly straight (stiff) filaments. **(C)** The filament assembly of desmin-p.S13P appears to be severely impaired. Only short fibrils with lengths from about 50 to a few hundred nanometers were observed. **(D)** Desmin-p.N107D shows coiled filaments and filamentous aggregates with a length up to a few micrometers. The small filament curvature suggests a rather low persistence length. **(E)** Desmin-p.N108G exhibits short filaments and protofilaments. The fringed hem of the filaments indicates that desmin monomers/oligomers are either only partially incorporated into the filament or that the filaments are inherently instable and easily decompose into their constituents. **(F)** Desmin-p.K109E shows short predominantly curly filamentous aggregates and protofilaments. The scale bars correspond to 1 µm or 200 nm **(E inset)** whereas the color bar corresponds to a height of 20 nm (WT, p.S13P, p.E108G), 30 nm (p.K109E) or 40nm (p.N107D), respectively.

Most of the known *DES* mutation carriers present a heterozygous genotype [22]. Subsequently, we performed co-transfection experiments using mutant desmin-EYFP constructs in combination with the red fluorescent wild-type desmin-mRuby construct to model a heterozygous status. Analysis of single transfected cells showed that there was no crosstalk between both fluorescent proteins under these experimental conditions (Supplements, Figure S6). In case of desmin-p.S13P, the majority of double transfected SW-13 cells showed filaments (Figure 6B) comparable to the wild-type double transfected cells (Figure 6A). Co-localization analysis determining the PCC [54] revealed that wild-type and desmin-p.S13P were incorporated into the same filaments similar to the wild-type / wild-type control (Figure 6A-B). In the majority of transfected SW-13 cells, desmin-p.N107D formed in the presence of wild-type desmin a mixture of filaments and cytoplasmic aggregates consisting of both desmin forms (Figure 6C-I). In contrast, desmin-p.E108G and -p.K109E dominantly inhibited the filament assembly if they were co-expressed together with wild-type desmin (Figure 6D-I). In H9c2 cells, we found similar effects of the investigated mutants, when co-expressed together with wild-type desmin (Figure 7). These findings might explain the dominant inheritance of desminopathies frequently observed in cardiovascular genetics.

**Figure 6.**
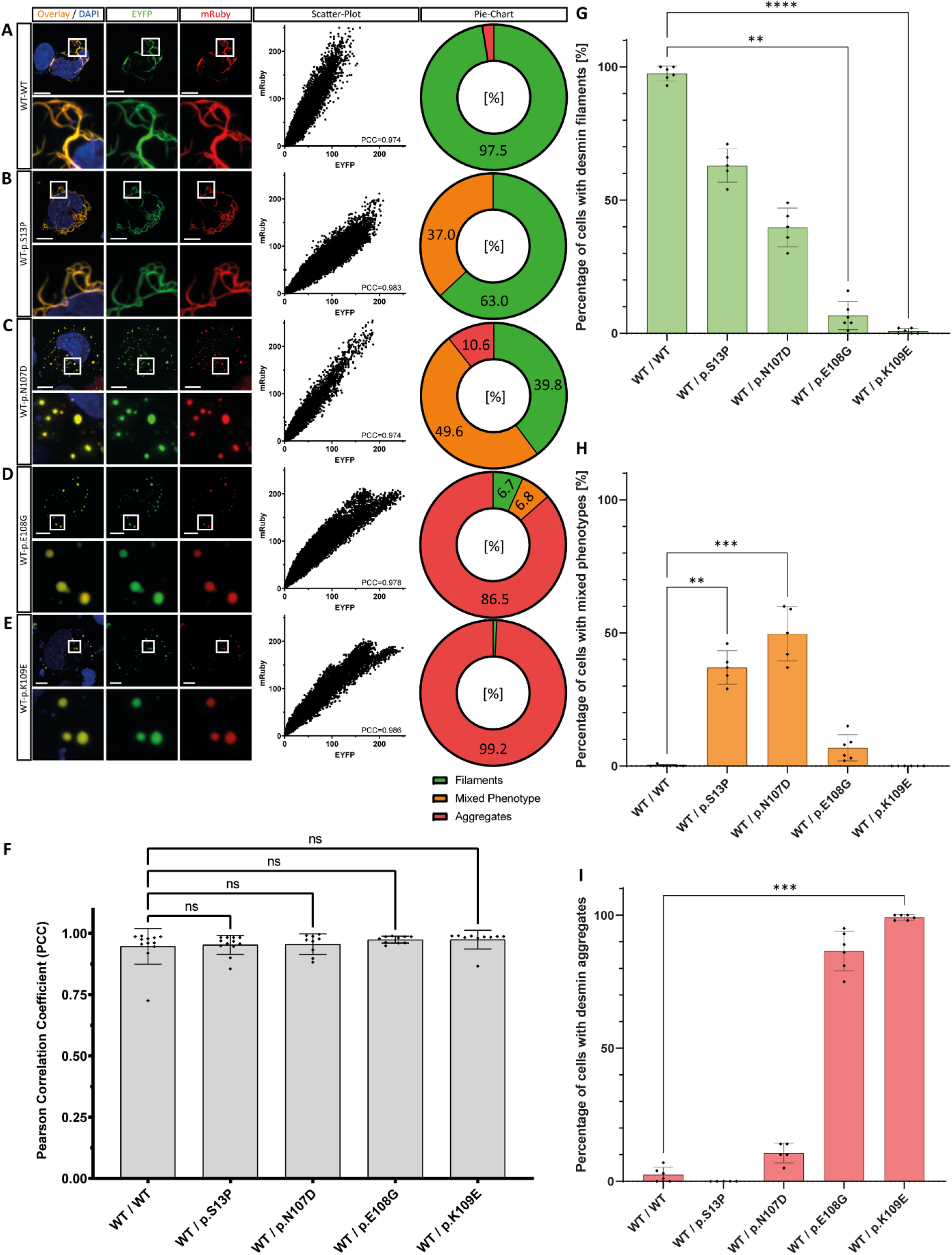
**(A-E)** Representative images of double transfected SW-13 cells, expressing wild-type desmin (fused to mRuby, red) and wild-type or mutant desmin (fused to EYFP, green). The overlay is shown in yellow. Nuclei were stained using DAPI (blue). Scale bars represent 10 µm. Representative scatter plots of the mRuby and EYFP channel used to determine the PCC are shown. The percentages of the different cell phenotypes are summarized as pie charts. **(F)** Statistical analysis of PCCs of the double transfected cells indicate a high colocalization of mutant and wild-type in filaments or aggregates. **(G-I)** Statistical analysis of desmin filament or aggregate formation using non-parametric Kruskal-Wallis test in double transfected SW-13 cells. All data are shown as mean ± SD. ns=not significant; *p≤0.05; **p≤0.01; ***p≤0.001; ****p≤0.0001.

**Figure 7.**
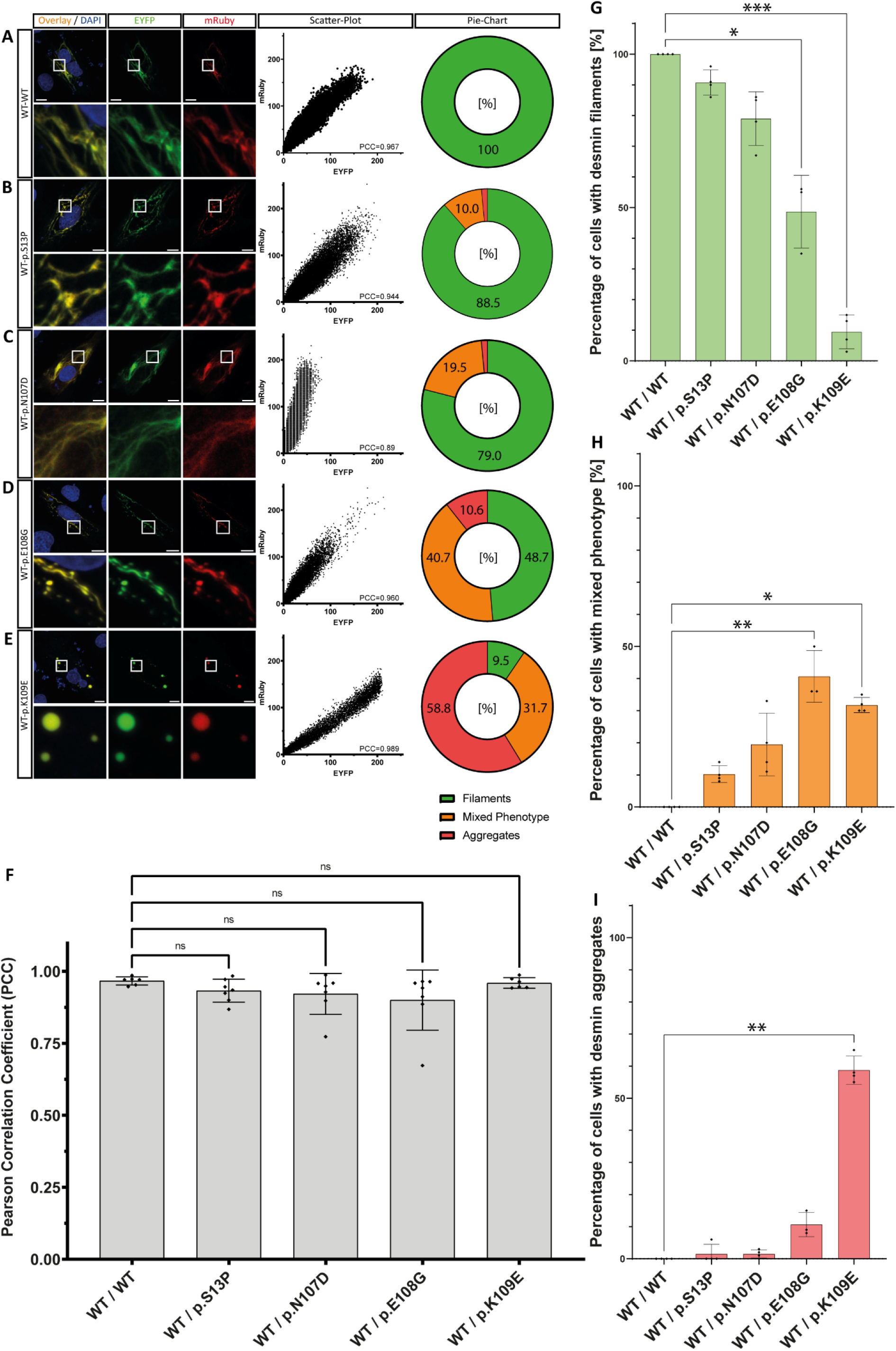
**(A-E)** Representative images of double transfected H9c2 cells, expressing wild-type desmin (fused mRuby, red) and wild-type or mutant desmin (fused to EYFP, green). The overlap is shown in yellow. Nuclei were stained using DAPI and are shown in blue. Scale bars represent 20 µm. Representative scatter plots of the mRuby and EYFP channel used to determine the PCC are shown. **(F)** Statistical analysis of PCCs of the double transfected cells indicate a high colocalization of mutant and wild-type in filaments or aggregates. **(G-I)** Statistical analysis of desmin filament or aggregate formation using non-parametric Kruskal-Wallis test in double transfected H9c2 cells. All data are shown as mean ± SD. *p≤0.05; **p≤0.01; ***p≤0.001; ****p≤0.0001.

Since it is known that the N-terminal head domain of desmin is necessary for the regular filament assembly [55], we systematically determined which parts of the head domain are necessary for filament formation by generation of deletion mutations (Figure 8A). In SW-13 cells, desmin mutants missing the first 10 or 20 amino acids behind the first methionine (p.S2-V11del and p.S2-A21del) were able to form similar filaments like the wild-type desmin (Figure 8B-D). All other larger N-terminal deletions caused a cytoplasmic aggregation or diffuse desmin localization in transfected SW-13 cells (Figure 8E-P). Interestingly, desmin deletion mutants missing 10 or 50 amino acids behind the first methionine were incorporated into filaments in the majority of transfected in H9c2 cells (Figure 9). However, larger deletions missing 60-110 amino acids behind the first methionine were unable to form filaments (Figure 9G-P).

**Figure 8.**
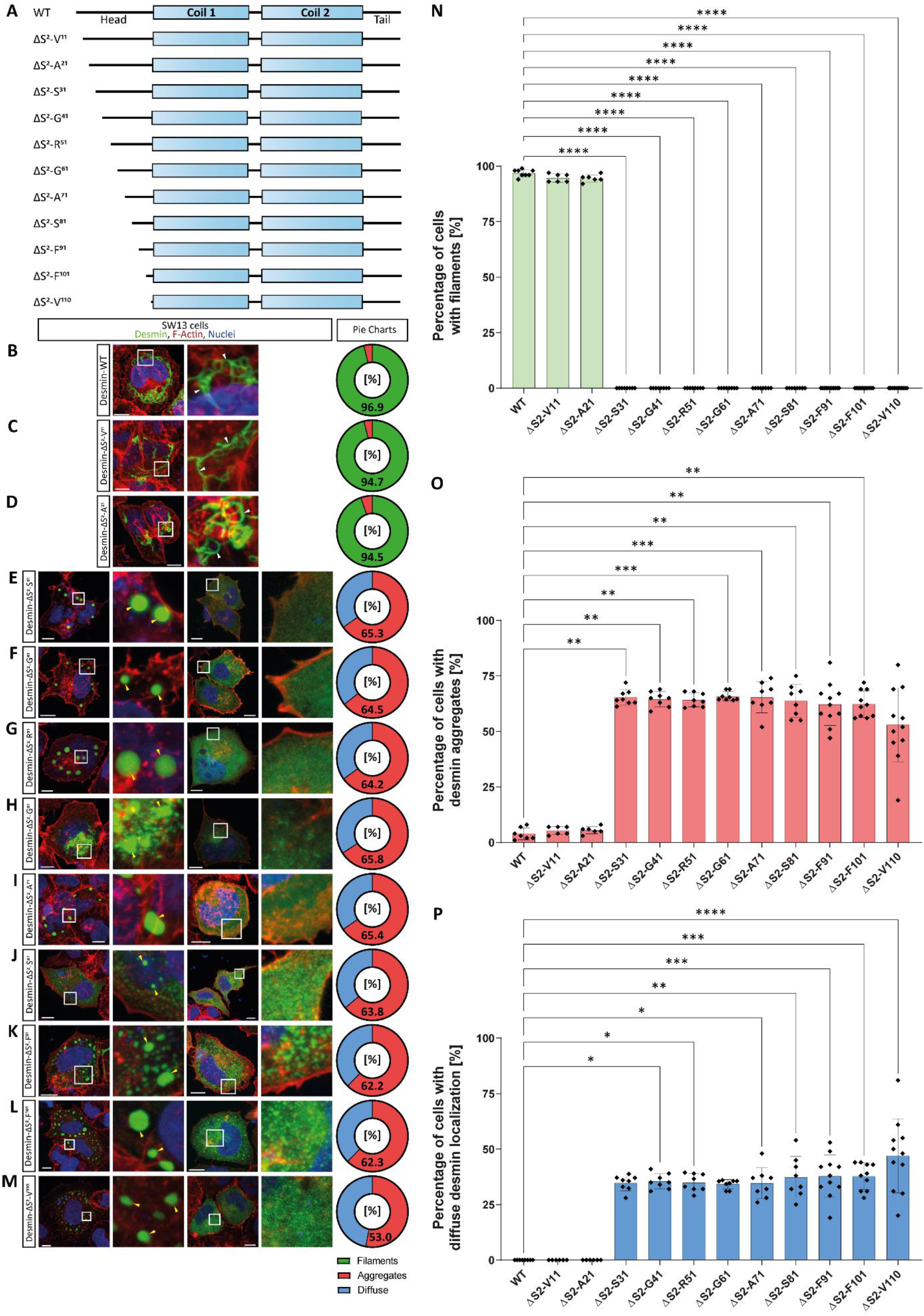
**(A)** Schematic overview about the generated N-terminal desmin deletion mutations. Representative images of SW-13 cells expressing wild-type desmin **(B)** or the deletion mutants **(C-M).** Desmin is shown in green, F-actin in red and the nuclei in blue. Scale bars represent 10 µm. Pie charts of observed subcellular desmin localization are shown **(B-M)**. Statistical analysis of desmin filament or aggregate formation was performed using non-parametric Kruskal-Wallis test in transfected SW-13 cells expressing wild-type desmin or desmin deletion mutants **(N-P)**. All data are shown as mean ± SD. *p≤0.05; **p≤0.01; ***p≤0.001; ****p≤0.0001.

**Figure 9.**
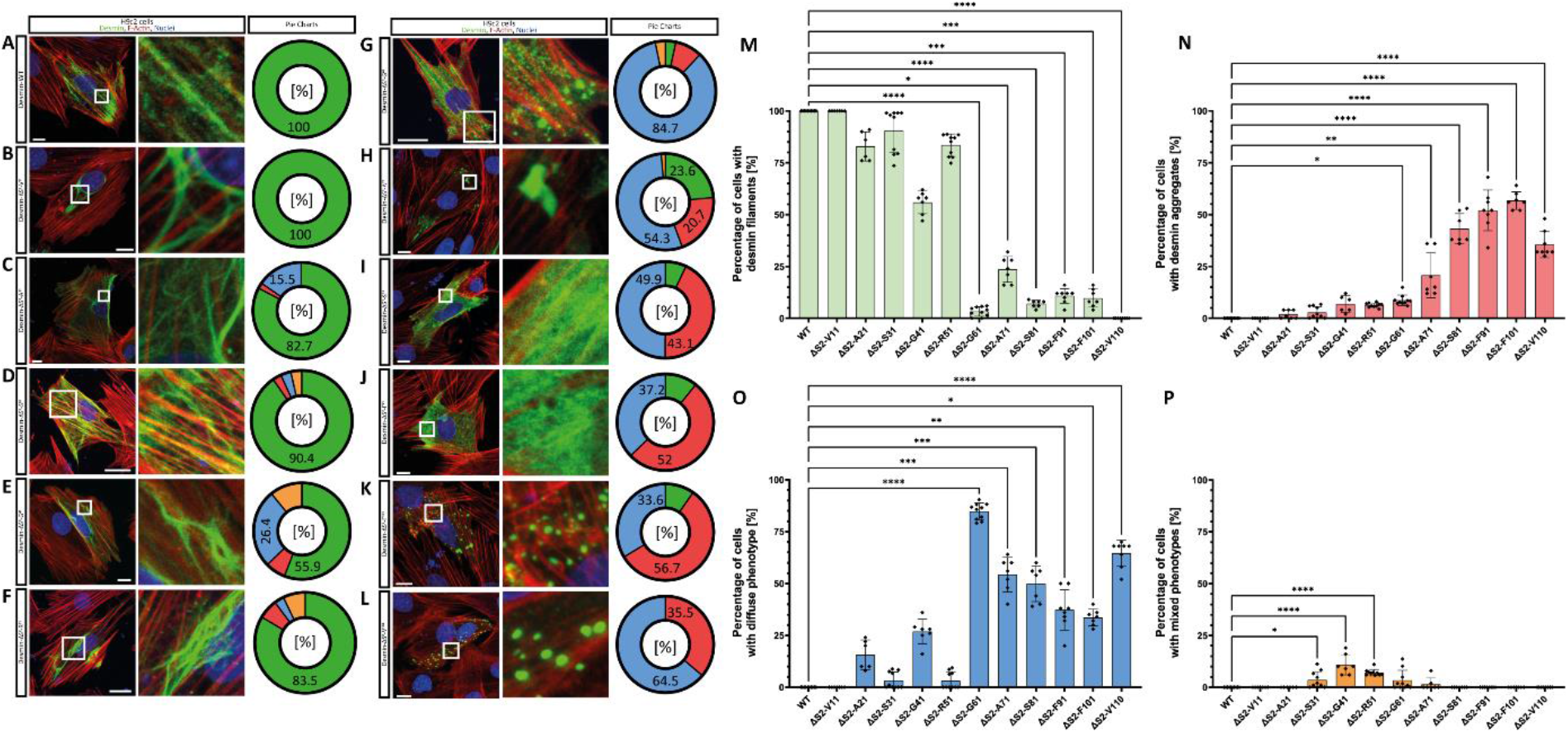
Representative images of SW-13 cells expressing wild-type desmin **(A)** or the deletion mutants **(B-L).** Desmin is shown in green, F-actin in red and the nuclei in blue. Scale bars represent 20 µm. Pie charts of observed cellular desmin localization are shown **(A-L)**. Statistical analysis of desmin filament or aggregate formation was performed using non-parametric Kruskal-Wallis test in transfected H9c2 cells expressing wild-type desmin or desmin deletion mutants **(M-P)**. All data are shown as mean ± SD. *p≤0.05; **p≤0.01; ***p≤0.001; ****p≤0.0001.

In summary, our study provides functional data about variants within the head domain of desmin and can be used in cardiovascular genetics as an ′atlas′ of *DES* variants contributing to variant classification.

## Discussion

Driven by a broad clinical application of next generation sequencing techniques and advanced classification criteria, the number of VUS significantly increased during the last years in cardiovascular genetics [56, 57]. However, VUS are frequently unsatisfying challenges for patients as well as for (cardiac) genetic counsellors. According to the guidelines of the American College of Medical Genetics and Genomics (ACMG) validated functional analysis of the mutant RNA or protein can be a strong criterion for variant classification [58, 59]. On the other hand, the majority of diagnostic laboratories is unable to perform functional analysis of VUS involved in cardiomyopathies, since dozens of genes with quite different cellular functions are involved [60]. One of these cardiomyopathy-associated genes is the *DES* gene encoding the muscle specific IF protein desmin, which provides cellular integrity of the (cardio)myocytes [22]. The most obvious hallmark of pathogenic *DES* mutations is an abnormal cytoplasmic aggregation and desmin accumulation [23, 27, 61]. However, the functional impact of the majority of rare *DES* variants listed in genetic disease databases is currently unknown. Therefore, we initiated this project investigating which of the known *DES* VUS affect the desmin filament formation *in vitro / in situ*.

Recently, we described a genetic hotspot in the 1A subdomain, where several likely pathogenic mutations affect the desmin filament formation [41]. Desmin consists of an N-terminal non-helical head domain [62], a central rod domain and a C-terminal tail domain [63]. The highly conserved rod domain mediates the coiled-coil formation and is divided into 1A, 1B and coil-2 subdomains [64]. Since the 1980s it is well known, that the non-helical head domain is necessary for filament assembly of type III IF proteins like desmin [55, 65]. In spite of this, the structural necessity of the head domain for filament assembly of type III IF proteins has been elucidated quite recently. Using cryo-electron microscopy and tomography Eibauer *et al.* showed that the head domains of vimentin molecules form an amyloid-like fiber in the center of the filament surrounded by five protofibrils [66]. Nevertheless, it is currently difficult to predict the impact of specific missense variants within the head domain on filament assembly. Our results indicate that the length of the N-terminal desmin head domain is highly relevant for the filament assembly. Systematic deletion of the N-terminus showed that small deletions of 20 amino acids did not affect the desmin filament assembly in SW-13 cells, whereas larger deletions with more than 30 deleted amino acids were unable to form IFs. Conversely, in H9c2 cells this effect started if more than 50 amino acids were deleted. The differences between these data sets might be explained by the endogenous desmin expression of H9c2 cells. Overall, these data are in good agreement with earlier reports by Kaufmann *et al.* using enzymatically truncated desmin missing the first 67 N-terminal amino acids [55]. Interestingly, synemin, an IF protein with a short N-terminal head domain, is also unable to form IFs. Synemin is incorporated into IFs, if it is co-expressed with other IF proteins like desmin or vimentin [67].

Clinically more relevant than deletions might be rare missense VUS, which are spread over the complete head domain of desmin (https://www.hgmd.cf.ac.uk/ and https://www.ncbi.nlm.nih.gov/clinvar/). Therefore, we inserted a set of 85 different *DES* missense VUS with a MAF<0.0001 which have been previously identified in patients with skeletal myopathies or cardiomyopathies and listed in disease databases. Using transiently transfected SW-13 cells in combination with confocal microscopy, we screened this set of VUS for filament assembly defects. By doing this, we identified four VUS (p.S13P, p.N107D, p.E108G and p.K109E), which caused an abnormal cytoplasmic desmin aggregation. Of note, *DES*-p.S13P and -p.E108G are localized at a position, where two different pathogenic *DES* mutations (p.S13F, p.E108K) have been previously described [15, 68, 69].

We verified these filaments defects likewise in H9c2 cells and in cardiomyocytes derived from iPSCs since these cells express endogenous desmin. Additionally, we investigated at the molecular level by *in vitro* assembly experiments in combination with AFM the molecular defects of these desmin mutants. The majority of missense mutations within the desmin head domain do not affect the filament formation (Figure 10). However, we can not exclude that nanomechanical properties of the mutant desmin filaments are affected as previously described by Kreplak *et al.* [49]. Of note, three of the investigated variants are localized at the C-terminus of the head domain in close proximity to the 1A-subdomain, which we recognized recently as a genetic hot spot indicating that this hot spot is extended. Recently, Eibauer *et al.* determined the structure of vimentin filaments and identified a molecular groove which they called interlock region [66]. Several amino acids, where mutations cause filament defects including p.N107D, p.E108G and p.K109E, are localized in this structural important interlock region.

**Figure 10.**
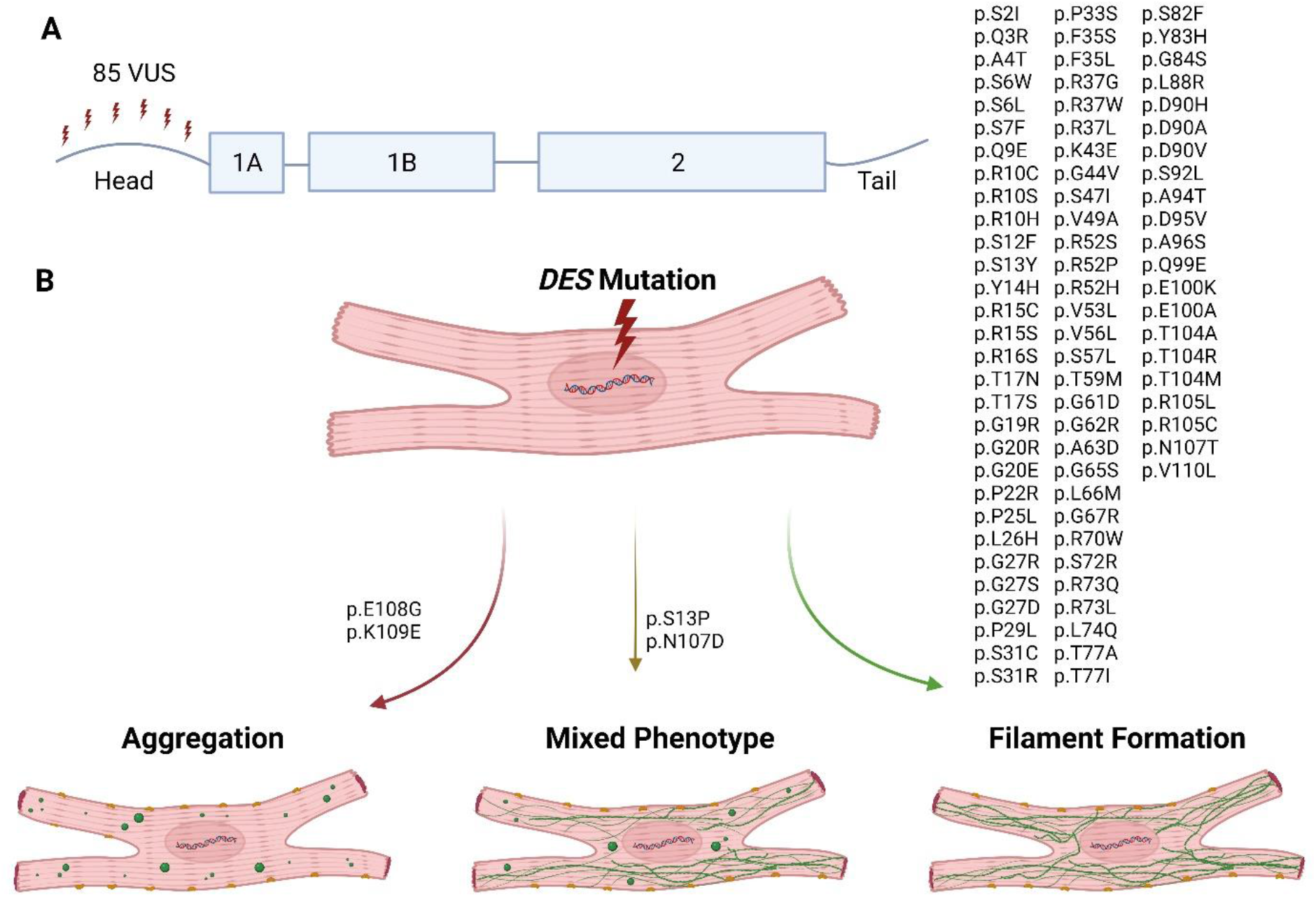
**(A)** A set of 85 different VUS localized in the desmin head domain have been investigated by cell transfection experiments in combination with confocal microscopy. **(B)** Schematic overview about the first part of the atlas of desmin (*DES*) mutations within the non-helical head domain.

Interestingly, the border region between head and 1A domain is likewise a genetic hot spot carrying pathogenic mutations in different IFs encoding genes like *GFAP* or *LMNA*, encoding glial fibrillary acidic protein or lamin A/C. *GFAP* mutations cause Alexander disease (OMIM, #203450; ORPHA:363717) [70] and *LMNA* mutations cause different cardiomyopathies or muscle dystrophies (OMIM, #115200; ORPHA:300751) [71–73]. Previously, different groups have identified pathogenic *GFAP* mutations at corresponding positions (p.E69K, p.R70Q and p.R70W) leading to Alexander disease [74–76]. Similarly, several pathogenic mutations associated with muscular dystrophies have been reported in the *LMNA* gene at corresponding positions (p.E31D, p.E31K, p.K32N, p.K32T) [77–79]. These reports in combination with our functional data about *DES* mutations highlight the general relevance of this region for filament assembly.

In consequence, clinicians and human geneticists should handle novel *DES* variants in the border region between the head and 1A domain with attention. Thus, several groups have identified disease-causing *DES* mutations in specific families [16, 20, 24, 80]. In conclusion, we present here the first part of a *DES* mutation ‘atlas’ (Figure 10) focussing on the functional characterization of VUS within the head domain, which can support disease classification and genetic counselling of affected mutation carriers.

## Acknowledgements

We thank Prof. Dr. Gerd Ulrich Nienhaus (Karlsruhe Institute of Technology, Germany) for providing the cDNA of mRuby [47]. In addition, we are thankful to Dr. Tomo Saric (University of Cologne, Germany) for providing the iPSC line NP00040-8 (UKKi011-A, https://ebisc.org/UKKi011-A/) generated from a healthy male donor.

