## Supplements for "Atlas of *DES* (desmin) variants: Impact of variants located within the head domain on filament assembly"

**Supplementary Figures**

**Figure S1.** Schematic maps of the used plasmids **(A)** pEYFP-N1-Desmin; **(B)** pmRuby-N1-Desmin and **(C)** pET100D-TOPO-Desmin.

**Figure S2.** Statistical analysis of aggregate and filament formation in transfected SW-13 cells. Statistical analysis of desmin filament or aggregate formation using non-parametric Kruskal-Wallis test in transfected SW-13 cells expressing wild-type desmin or desmin deletion mutants. All data are shown as mean ± standard deviation. *p≤0.05; **p≤0.01; ***p≤0.001; ****p≤0.0001.

**Figure S3.** Expression analysis. **(A)** SW-13 and H9c2 cells were transfected with desmin constructs fused to EYFP. Fluorescence was analysed using a plate reader. Statistical analysis of the normalized fluorescence intensities revealed no obvious differences between wild-type and mutant desmin in **(B)** SW-13 and **(C)** H9c2 cells.

**Figure S4.** Purification of recombinant desmin by ionic exchange chromatography (IEC). **(A)** SDS-PAGE in combination with Coomassie-R250 staining of the fractions of the IEC. The elution fractions (8-10) were pooled and used afterwards for IMAC. **(B)** Chromatogram showing the elution of recombinant desmin from the IEC column. A linear gradient with increasing [NaCl] was used for elution.

**Figure S5.** Purification of recombinant desmin by immobilized metal affinity chromatography (IMAC). **(A)** SDS-PAGE in combination with Coomassie-R250 staining of the fractions of the IMAC. **(B)** Chromatogram showing the elution of recombinant desmin from the HisTrap column. A stepwise gradient with increased imidazole concentration was used for elution.

**Figure S6.** Fluorescence analysis in single transfected SW-13 cells. Representative images of single transfected cells expressing wild-type desmin fused with **(A)** mRuby or **(B)** EYFP. Scale bars represent 10 µm. Scatter plots indicate an absence of a significant crosstalk between both fluorescent proteins.

**Figure S1**

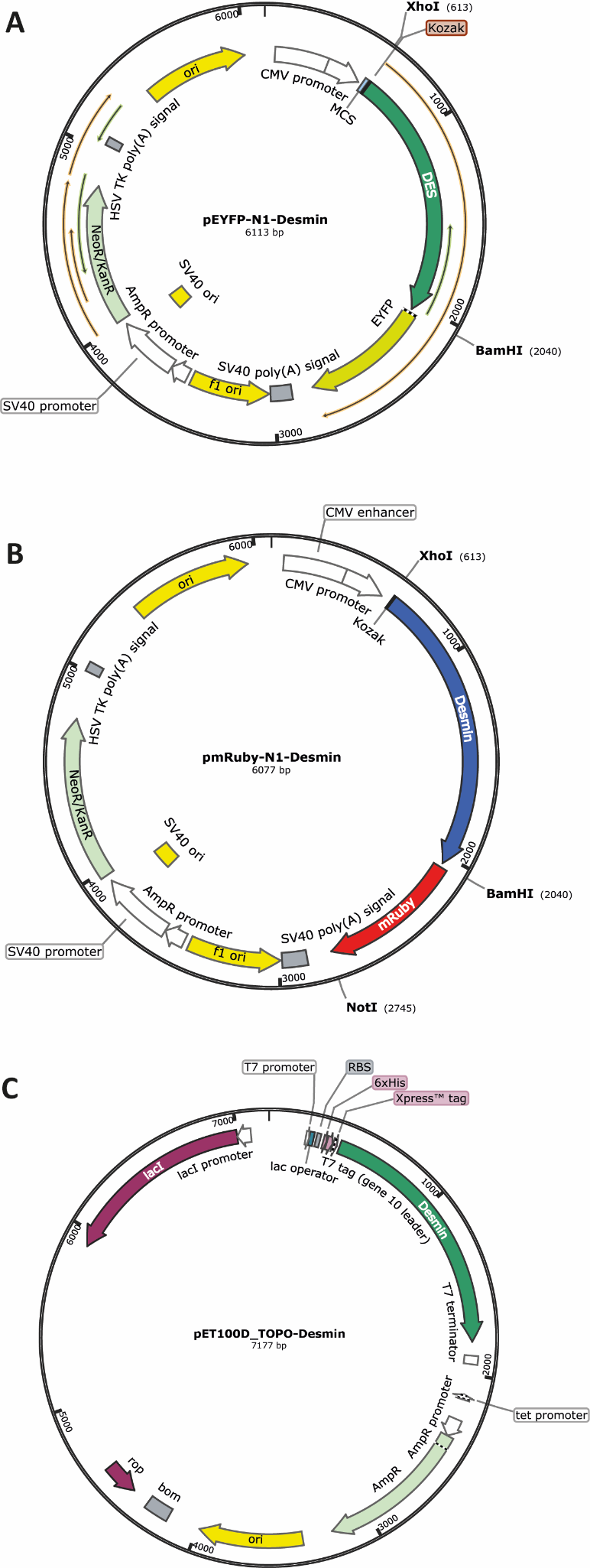

**Figure S2**

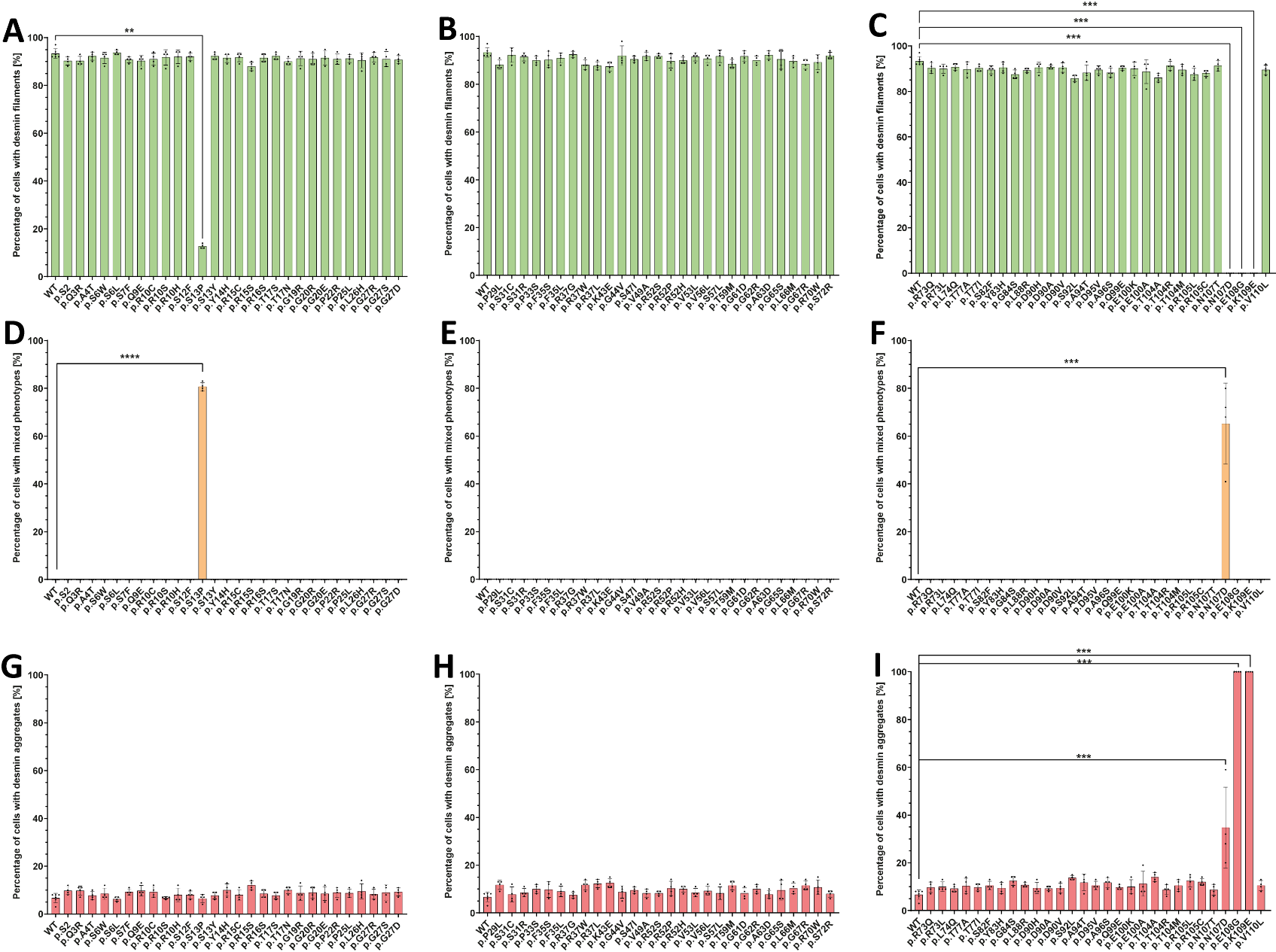

**Figure S3**

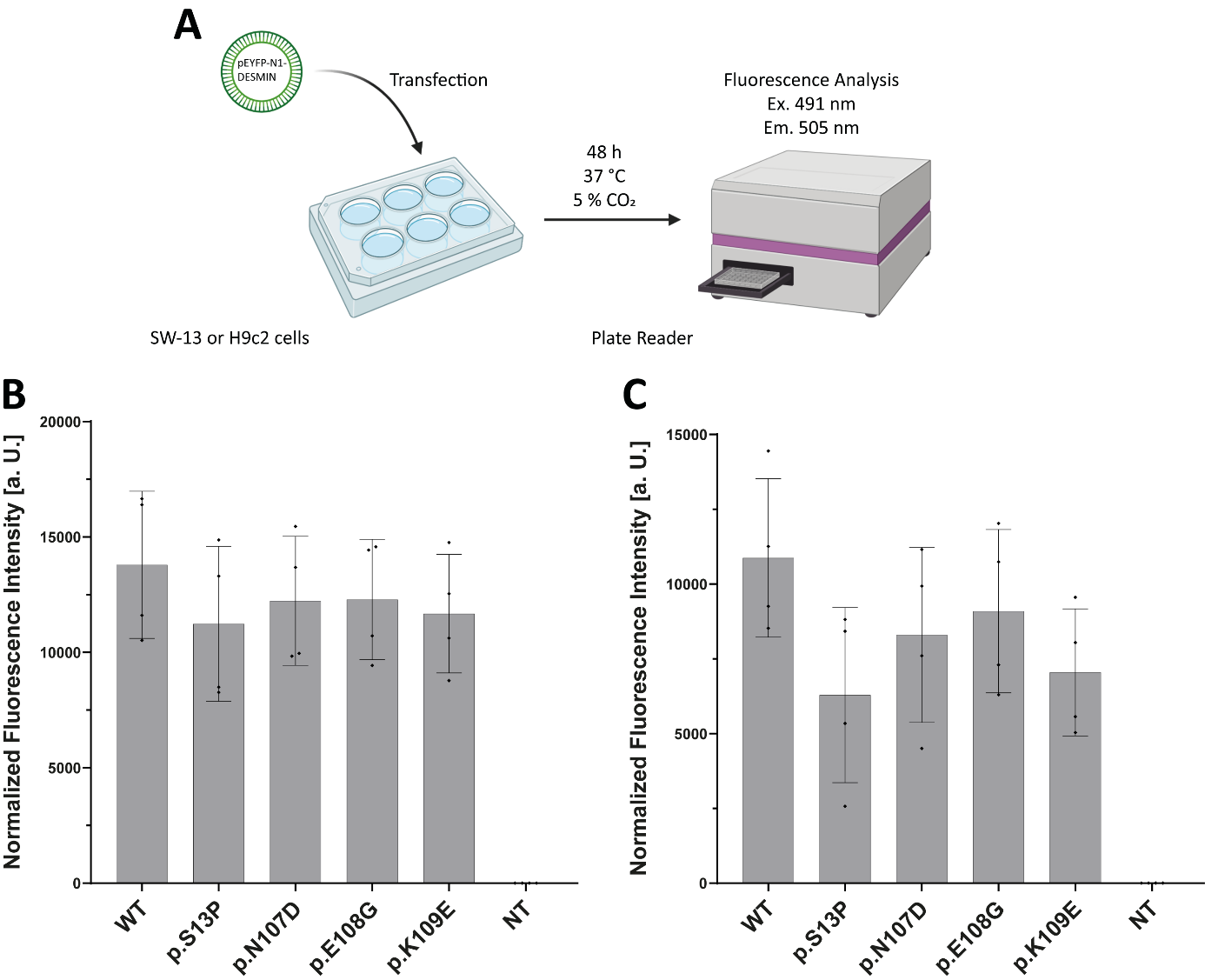

**Figure S4**

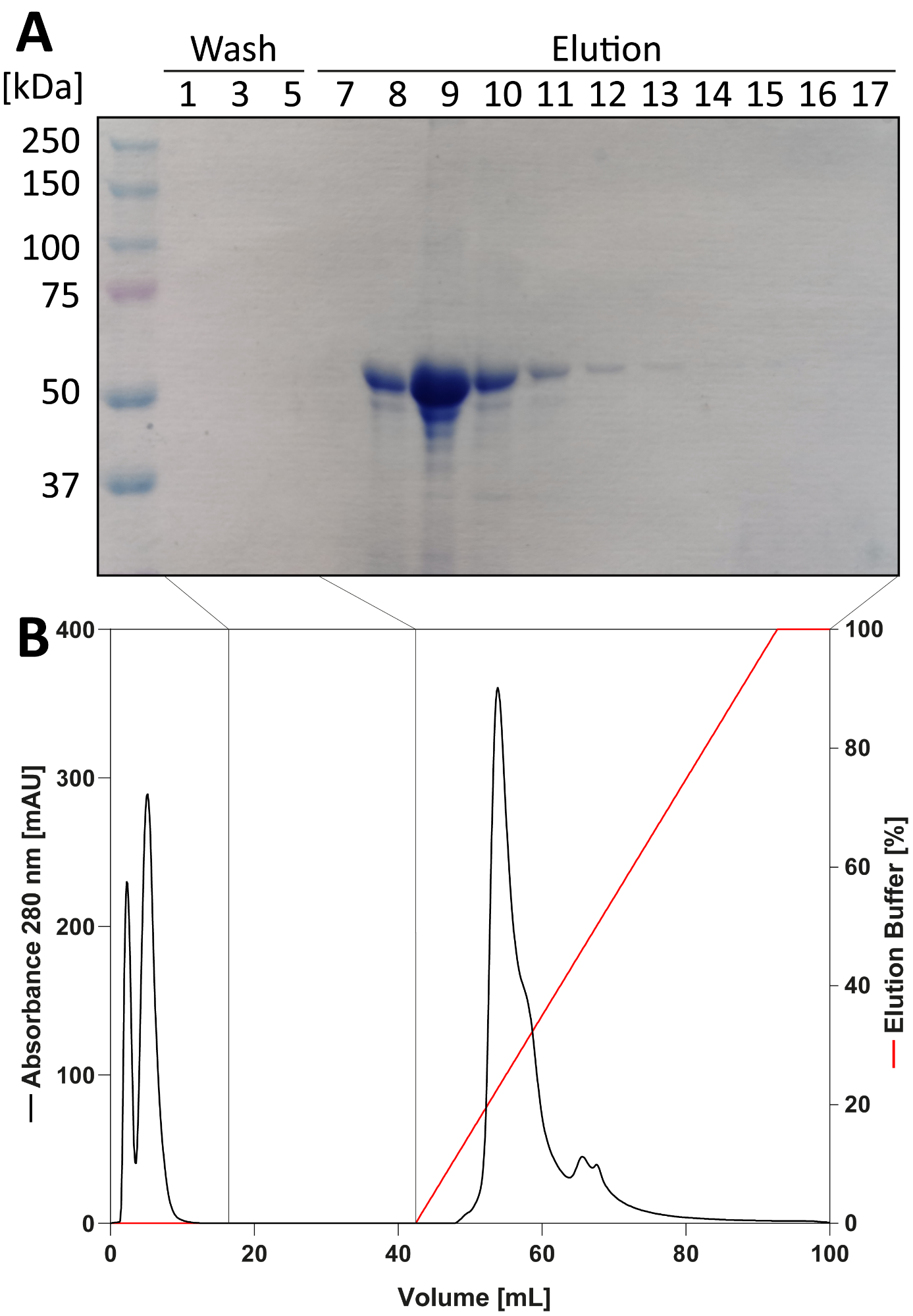

**Figure S5**

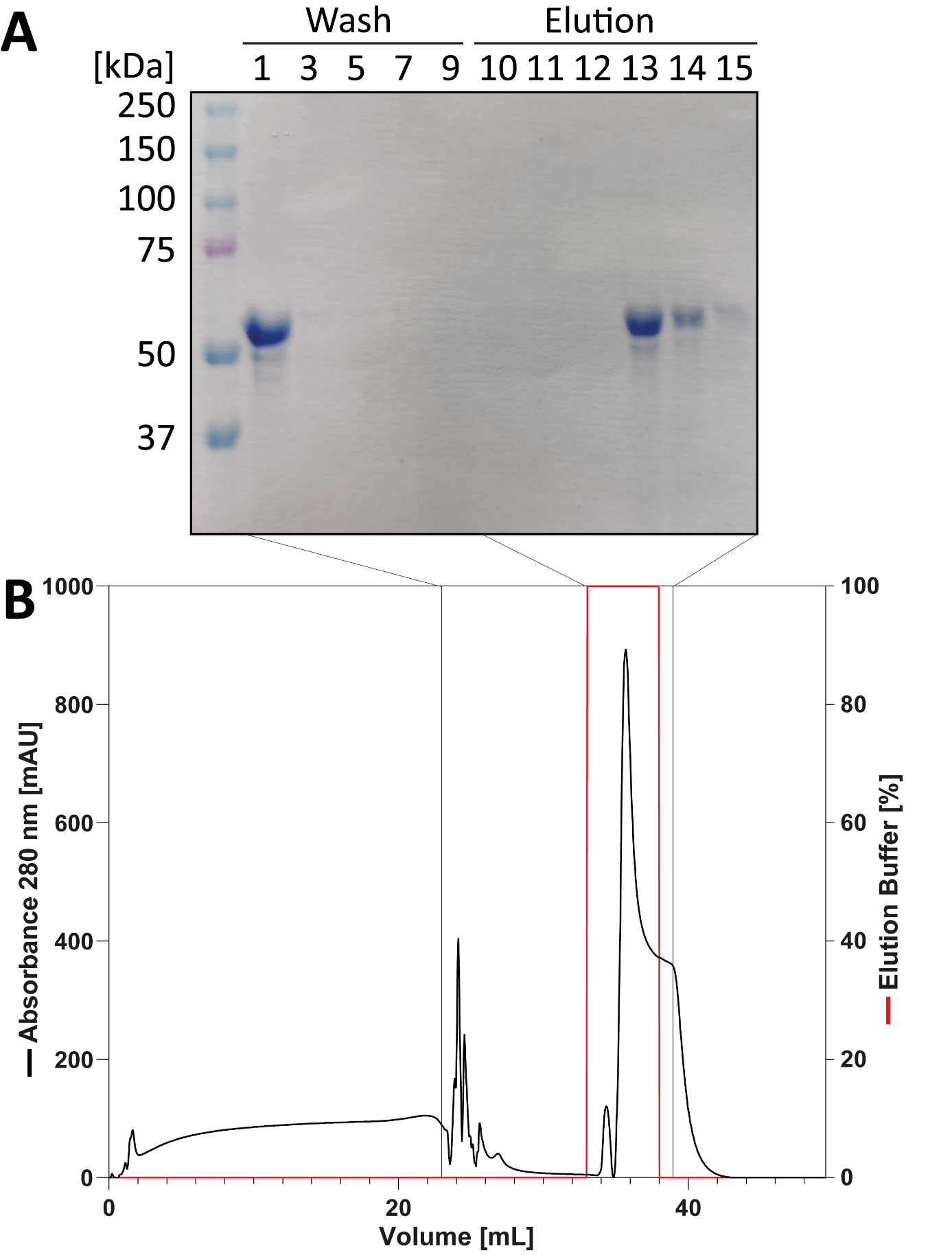

**Figure S6**

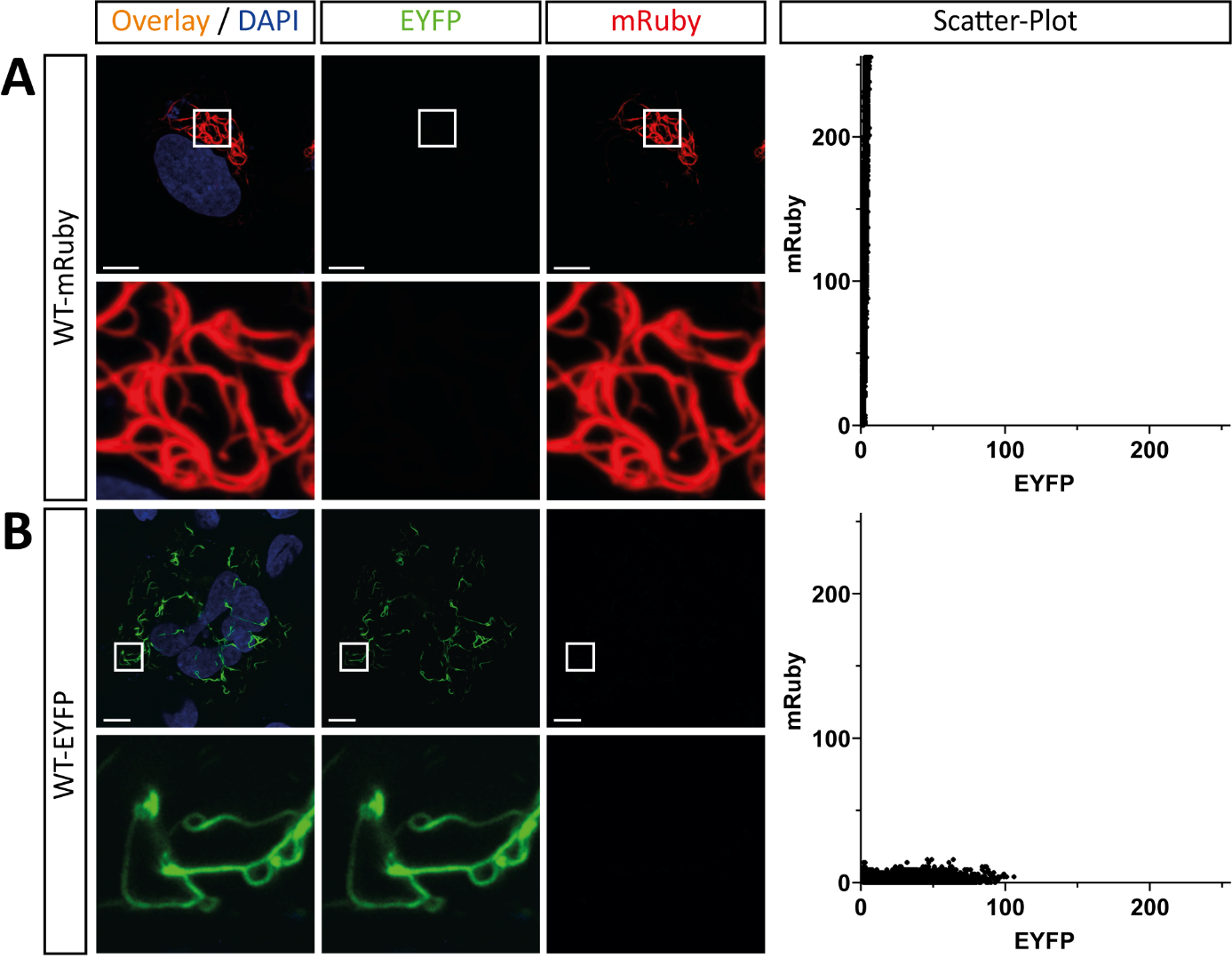

**Table S1.** Overview about the used oligonucleotides.

| Name | Sequence 5‘-3‘ | Application |
| --- | --- | --- |
| DES_L110V_for | GCACCAACGAGAAG**T**TGGAGCTGCAGGAG | SDM |
| DES_L110V_rev | CTCCTGCAGCTCCA**A**CTTCTCGTTGGTGC | SDM |
| DES_K109E_for | GCGCACCAACGAG**G**AGGTGGAGCTGCA | SDM |
| DES_K109E_rev | TGCAGCTCCACCT**C**CTCGTTGGTGCGC | SDM |
| DES_E108G_for | ACGCGCACCAACG**G**GAAGGTGGAGCTG | SDM |
| DES_E108G_rev | CAGCTCCACCTTC**C**CGTTGGTGCGCGT | SDM |
| DES_N107D_for | GACCACGCGCACC**G**ACGAGAAGGTGGA | SDM |
| DES_N107D_rev | TCCACCTTCTCGT**C**GGTGCGCGTGGTC | SDM |
| DES_N107T_for | GACCACGCGCA**C**CACCGAGAAGGTGGA | SDM |
| DES_N107T_rev | TCCACCTTCTCG**G**TGGTGCGCGTGGTC | SDM |
| DES_R105L_for | GAGTTTCTGACCACGC**T**CACCAACGAGAAGGTG | SDM |
| DES_R105L_rev | CACCTTCTCGTTGGTG**A**GCGTGGTCAGAAACTC | SDM |
| DES_R105C_for | GAGTTTCTGACCACG**T**GCACCAACGAGAAGG | SDM |
| DES_R105C_rev | CCTTCTCGTTGGTGC**A**CGTGGTCAGAAACTC | SDM |
| DES_T104M_for | CCAGGAGTTTCTGACCA**T**GCGCACCAACG | SDM |
| DES_T104M_rev | CGTTGGTGCGC**A**TGGTCAGAAACTCCTGG | SDM |
| DES_T104R_for | CCAGGAGTTTCTGACCA**G**GCGCACCAACG | SDM |
| DES_T104R_rev | CGTTGGTGCGC**C**TGGTCAGAAACTCCTGG | SDM |
| DES_T104A_for | AGGAGTTTCTGACC**G**CGCGCACCAACGAG | SDM |
| DES_T104A_rev | CTCGTTGGTGCGCG**C**GGTCAGAAACTCCT | SDM |
| DES_E100A_for | CGCGGTGAACCAGG**C**GTTTCTGACCACGC | SDM |
| DES_E100A_rev | GCGTGGTCAGAAAC**G**CCTGGTTCACCGCG | SDM |
| DES_E100K_for | GACGCGGTGAACCAG**A**AGTTTCTGACCACGC | SDM |
| DES_E100K_rev | GCGTGGTCAGAAACT**T**CTGGTTCACCGCGTC | SDM |
| DES_Q99E_for | CCGACGCGGTGAAC**G**AGGAGTTTCTGACC | SDM |
| DES_Q99E_rev | GGTCAGAAACTCCT**C**GTTCACCGCGTCGG | SDM |
| DES_A96S_for | CTCACTGGCCGAC**T**CGGTGAACCAGGA | SDM |
| DES_A96S_rev | TCCTGGTTCACCG**A**GTCGGCCAGTGAG | SDM |
| DES_D95V_for | TTCTCACTGGCCG**T**CGCGGTGAACCAG | SDM |
| DES_D95V_rev | CTGGTTCACCGCG**A**CGGCCAGTGAGAA | SDM |
| DES_A94T_for | TGGACTTCTCACTG**A**CCGACGCGGTGAAC | SDM |
| DES_A94T_rev | GTTCACCGCGTCGG**T**CAGTGAGAAGTCCA | SDM |
| DES_S92L_for | GGCGAGCTGCTGGACTTCT**T**ACTGGCCGAC | SDM |
| DES_S92L_rev | GTCGGCCAGT**A**AGAAGTCCAGCAGCTCGCC | SDM |
| DES_D90V_for | GGCGAGCTGCTGG**T**CTTCTCACTGGCC | SDM |
| DES_D90V_rev | GGCCAGTGAGAAG**A**CCAGCAGCTCGCC | SDM |
| DES_D90A_for | GGCGAGCTGCTGG**C**CTTCTCACTGGCC | SDM |
| DES_D90A_rev | GGCCAGTGAGAAG**G**CCAGCAGCTCGCC | SDM |
| DES_D90H_for | AGGCGAGCTGCTG**C**ACTTCTCACTGGC | SDM |
| DES_D90H_rev | GCCAGTGAGAAGT**G**CAGCAGCTCGCCT | SDM |
| DES_L88R_for | GCGCAGGCGAGC**G**GCTGGACTTCTC | SDM |
| DES_L88R_rev | GAGAAGTCCAGC**C**GCTCGCCTGCGC | SDM |
| DES_G84S_for | CCCTCCTCCTAC**A**GCGCAGGCGAGC | SDM |
| DES_G84S_rev | GCTCGCCTGCGC**T**GTAGGAGGAGGG | SDM |
| DES_Y83H_for | CGCCCTCCTCC**C**ACGGCGCAGGC | SDM |
| DES_Y83H_rev | GCCTGCGCCGT**G**GGAGGAGGGCG | SDM |
| DES_S82F_for | GCACGCCCTCCT**T**CTACGGCGCAGG | SDM |
| DES_S82F_rev | CCTGCGCCGTAG**A**AGGAGGGCGTGC | SDM |
| DES_T77I_for | CCGGCTGGGGACCA**T**CCGCACGC | SDM |
| DES_T77I_rev | GCGTGCGG**A**TGGTCCCCAGCCGG | SDM |
| DES_T77A_for | GGCTGGGGACC**G**CCCGCACGCCC | SDM |
| DES_T77A_rev | GGGCGTGCGGG**C**GGTCCCCAGCC | SDM |
| DES_L74Q_for | GGCCAGCCGGC**A**GGGGACCACCC | SDM |
| DES_L74Q_rev | GGGTGGTCCCC**T**GCCGGCTGGCC | SDM |
| DES_R73L_for | GCGGGCCAGCC**T**GCTGGGGACCA | SDM |
| DES_R73L_rev | TGGTCCCCAGC**A**GGCTGGCCCGC | SDM |
| DES_R73Q_for | GCGGGCCAGCC**A**GCTGGGGACCA | SDM |
| DES_R73Q_rev | TGGTCCCCAGC**T**GGCTGGCCCGC | SDM |
| DES_S72R_for | CTGCGGGCCAG**G**CGGCTGGGGAC | SDM |
| DES_S72R_rev | GTCCCCAGCCG**C**CTGGCCCGCAG | SDM |
| DES_R70W_for | TGGGGTCGCTG**T**GGGCCAGCCGG | SDM |
| DES_R70W_rev | CCGGCTGGCCC**A**CAGCGACCCCA | SDM |
| DES_G67R_for | CCGGGGGCCTG**A**GGTCGCTGCGG | SDM |
| DES_G67R_rev | CCGCAGCGACC**T**CAGGCCCCCGG | SDM |
| DES_L66M_for | GGGCCGGGGGC**A**TGGGGTCGCTG | SDM |
| DES_L66M_rev | CAGCGACCCCA**T**GCCCCCGGCCC | SDM |
| DES_G65S_for | CGGGGCCGGG**A**GCCTGGGGTC | SDM |
| DES_G65S_rev | GACCCCAGGC**T**CCCGGCCCCG | SDM |
| DES_A63D_for | TCGGGCGGGG**A**CGGGGGCCTG | SDM |
| DES_A63D_rev | CAGGCCCCCG**T**CCCCGCCCGA | SDM |
| DES_G62R_for | CACGTCGGGC**A**GGGCCGGGGG | SDM |
| DES_G62R_rev | CCCCCGGCCC**T**GCCCGACGTG | SDM |
| DES_G61D_for | CGCACGTCGG**A**CGGGGCCGGG | SDM |
| DES_G61D_rev | CCCGGCCCCG**T**CCGACGTGCG | SDM |
| DES_T59M_for | GGTGTCGCGCA**T**GTCGGGCGGGG | SDM |
| DES_T59M_rev | CCCCGCCCGAC**A**TGCGCGACACC | SDM |
| DES_S57L_for | CCGCGTGTACCAGGTGT**T**GCGCACGTC | SDM |
| DES_S57L_rev | GACGTGCGC**A**ACACCTGGTACACGCGG | SDM |
| DES_V56L_for | CCGCGTGTACCAG**T**TGTCGCGCACGTC | SDM |
| DES_V56L_rev | GACGTGCGCGAC**A**ACTGGTACACGCGG | SDM |
| DES_V53L_for | GGTGACGTCCCGC**T**TGTACCAGGTGTC | SDM |
| DES_V53L_rev | GACACCTGGTAC**A**AGCGGGACGTCACC | SDM |
| DES_R52H_for | TCGGTGACGTCCC**A**CGTGTACCAGGTG | SDM |
| DES_R52H_rev | CACCTGGTACACG**T**GGGACGTCACCGA | SDM |
| DES_R52P_for | CGGTGACGTCCC**C**CGTGTACCAGGT | SDM |
| DES_R52P_rev | ACCTGGTACACG**G**GGGACGTCACCG | SDM |
| DES_R52S_for | CTCGGTGACGTCC**A**GCGTGTACCAGGT | SDM |
| DES_R52S_rev | ACCTGGTACACGC**T**GGACGTCACCGAG | SDM |
| DES_V49A_for | CTCCAGCTCGG**C**GACGTCCCGCG | SDM |
| DES_V49A_rev | CGCGGGACGTC**G**CCGAGCTGGAG | SDM |
| DES_S47I_for | AAGGGCTCCTCCA**T**CTCGGTGACGTCC | SDM |
| DES_S47I_rev | GGACGTCACCGAG**A**TGGAGGAGCCCTT | SDM |
| DES_G44V_for | TTTCGGCTCTAAGG**T**CTCCTCCAGCTCGG | SDM |
| DES_G44V_rev | CCGAGCTGGAGGAG**A**CCTTAGAGCCGAAA | SDM |
| DES_K43E_for | GGGCGGGTTTCGGCTCT**G**AGGGCTCCT | SDM |
| DES_K43E_rev | AGGAGCCCT**C**AGAGCCGAAACCCGCCC | SDM |
| DES_R37L_for | CCGTGTTCCCGC**T**GGCGGGTTTCG | SDM |
| DES_R37L_rev | CCGAAACCCGCC**A**GCGGGAACACGG | SDM |
| DES_R37W_for | CCCGTGTTCCCG**T**GGGCGGGTTTCG | SDM |
| DES_R37W_rev | CGAAACCCGCCC**A**CGGGAACACGGG | SDM |
| DES_R37G_for | CCGTGTTCCCG**G**GGGCGGGTTTC | SDM |
| DES_R37G_rev | GAAACCCGCCC**C**CGGGAACACGG | SDM |
| DES_F35L_for | GAGCTCGCCCGTGTT**A**CCGCGGGC | SDM |
| DES_F35L_rev | GCCCGCGG**T**AACACGGGCGAGCTC | SDM |
| DES_F35S_for | GCTCGCCCGTG**AG**CCCGCGGGCGG | SDM |
| DES_F35S_rev | CCGCCCGCGGG**CT**CACGGGCGAGC | SDM |
| DES_P33S_for | CCCGCTGAGCTCG**AG**CGTGTTCCCGCGG | SDM |
| DES_P33S_rev | CCGCGGGAACACG**CT**CGAGCTCAGCGGG | SDM |
| DES_S31R_for | CTCCCCGCTGAG**G**TCGCCCGTGTTC | SDM |
| DES_S31R_rev | GAACACGGGCGA**C**CTCAGCGGGGAG | SDM |
| DES_S31C_for | GCTCCCCGCTG**T**GCTCGCCCGTG | SDM |
| DES_S31C_rev | CACGGGCGAGC**A**CAGCGGGGAGC | SDM |
| DES_P29L_for | GCTCGGCTCCC**T**GCTGAGCTCGC | SDM |
| DES_P29L_rev | GCGAGCTCAGC**A**GGGAGCCGAGC | SDM |
| DES_G27D_for | GCTTCCCGCTCG**A**CTCCCCGCTGAG | SDM |
| DES_G27D_rev | CTCAGCGGGGAG**T**CGAGCGGGAAGC | SDM |
| DES_G27S_for | GCTTCCCGCTC**A**GCTCCCCGCTG | SDM |
| DES_G27S_rev | CAGCGGGGAGC**T**GAGCGGGAAGC | SDM |
| DES_G27R_for | GCTTCCCGCTC**C**GCTCCCCGCTG | SDM |
| DES_G27R_rev | CAGCGGGGAGC**G**GAGCGGGAAGC | SDM |
| DES_L26H_for | GGGCTTCCCGC**A**CGGCTCCCCGC | SDM |
| DES_L26H_rev | GCGGGGAGCCG**T**GCGGGAAGCCC | SDM |
| DES_P25L_for | CCCGGGCTTCC**T**GCTCGGCTCCC | SDM |
| DES_P25L_rev | GGGAGCCGAGC**A**GGAAGCCCGGG | SDM |
| DES_P22R_for | GGCGGGGCCC**G**GGGCTTCCCG | SDM |
| DES_P22R_rev | CGGGAAGCCC**C**GGGCCCCGCC | SDM |
| DES_G20E_for | CACCTTCGGCG**A**GGCCCCGGGCT | SDM |
| DES_G20E_rev | AGCCCGGGGCC**T**CGCCGAAGGTG | SDM |
| DES_G20R_for | GCACCTTCGGC**A**GGGCCCCGGGC | SDM |
| DES_G20R_rev | GCCCGGGGCCC**T**GCCGAAGGTGC | SDM |
| DES_G19R_for | CCGCACCTTC**C**GCGGGGCCCC | SDM |
| DES_G19R_rev | GGGGCCCCGC**G**GAAGGTGCGG | SDM |
| DES_T17N_for | CTACCGCCGCA**A**CTTCGGCGGGG | SDM |
| DES_T17N_rev | CCCCGCCGAAGT**T**GCGGCGGTAG | SDM |
| DES_T17S_for | CTACCGCCGCA**G**CTTCGGCGGGG | SDM |
| DES_T17S_rev | CCCCGCCGAAG**C**TGCGGCGGTAG | SDM |
| DES_R16S_for | TCCTCCTACCGC**A**GCACCTTCGGCG | SDM |
| DES_R16S_rev | CGCCGAAGGTGC**T**GCGGTAGGAGGA | SDM |
| DES_R15S_for | CGTGTCCTCCTAC**A**GCCGCACCTTCGG | SDM |
| DES_R15S_rev | CCGAAGGTGCGGC**T**GTAGGAGGACACG | SDM |
| DES_R15C_for | CGTGTCCTCCTAC**T**GCCGCACCTTCGG | SDM |
| DES_R15C_rev | CCGAAGGTGCGGC**A**GTAGGAGGACACG | SDM |
| DES_Y14H_for | GCGTGTCCTCCC**A**CCGCCGCACC | SDM |
| DES_Y14H_rev | GGTGCGGCGGT**G**GGAGGACACGC | SDM |
| DES_S13Y_for | CCAGCGCGTGTCCT**AT**TACCGCCGCACCTT | SDM |
| DES_S13Y_rev | AAGGTGCGGCGGTA**AT**AGGACACGCGCTGG | SDM |
| DES_S13P_for | AGCGCGTGTCC**C**CCTACCGCCGC | SDM |
| DES_S13P_rev | GCGGCGGTAGG**G**GGACACGCGCT | SDM |
| DES_S12F_for | GCCAGCGCGTGT**T**CTCCTACCGCCG | SDM |
| DES_S12F_rev | CGGCGGTAGGAG**A**ACACGCGCTGGC | SDM |
| DES_R10H_for | TCGTCCAGCCAGC**A**CGTGTCCTCCTAC | SDM |
| DES_R10H_rev | GTAGGAGGACACG**T**GCTGGCTGGACGA | SDM |
| DES_R10S_for | CTCGTCCAGCCAG**A**GCGTGTCCTCCTA | SDM |
| DES_R10S_rev | TAGGAGGACACGC**T**CTGGCTGGACGAG | SDM |
| DES_R10C_for | CTCGTCCAGCCAG**T**GCGTGTCCTCCTA | SDM |
| DES_R10C_rev | TAGGAGGACACGC**A**CTGGCTGGACGAG | SDM |
| DES_Q9E_for | CTACTCGTCCAGC**G**AGCGCGTGTCCTC | SDM |
| DES_Q9E_rev | GAGGACACGCGCT**C**GCTGGACGAGTAG | SDM |
| DES_S7F_for | CAGGCCTACTCGT**T**CAGCCAGCGCGTG | SDM |
| DES_S7F_rev | CACGCGCTGGCTG**A**ACGAGTAGGCCTG | SDM |
| DES_S6L_for | GCCAGGCCTACT**T**GTCCAGCCAGCG | SDM |
| DES_S6L_rev | CGCTGGCTGGAC**A**AGTAGGCCTGGC | SDM |
| DES_S6W_for | GCCAGGCCTACT**G**GTCCAGCCAGCG | SDM |
| DES_S6W_rev | CGCTGGCTGGAC**C**AGTAGGCCTGGC | SDM |
| DES_A4T_for | TCACCATGAGCCAG**A**CCTACTCGTCCAGC | SDM |
| DES_A4T_rev | GCTGGACGAGTAGGTC**T**GGCTCATGGTGA | SDM |
| DES_Q3R_for | GTCACCATGAGCC**G**GGCCTACTCGTCC | SDM |
| DES_Q3R_rev | GGACGAGTAGG**C**CCGGCTCATGGTGAC | SDM |
| DES_S2I_for | CGAGGCCGTCACCATGA**T**CCAGGCCTA | SDM |
| DES_S2I_rev | TAGGCCTGG**A**TCATGGTGACGGCCTCG | SDM |
| CMV_for | CGCAAATGGGCGGTAGGCGTG | Sanger Sequencing |
| EGFP_N_rev | GCTTGCCGTAGGTGGCATC | Sanger Sequencing |
| T7_for | TAATACGACTCACTATAGGG | Sanger Sequencing |
| T7_rev | GCTAGTTATTGCTCAGCGGT | Sanger Sequencing |
| DES_HEAD_del_for | GGCCGTCACCATGGAGCTGCAGGAGC | SDM |
| DES_HEAD_del_rev | GCTCCTGCAGCTCCATGGTGACGGCC | SDM |
| DES-S2-V11del_for | GGCCGTCACCATGTCCTCCTACCGCC | SDM |
| DES-S2-V11del_rev | GGCGGTAGGAGGACATGGTGACGGCC | SDM |
| DES-S2-A21del_for | CGGGAAGCCCGGCATGGTGACGGC | SDM |
| DES-S2-A21del_rev | GCCGTCACCATGCCGGGCTTCCCG | SDM |
| DES-S2-G41del_for | GGAGGAGCCCTTAGACATGGTGACGGCCTC | SDM |
| DES-S2-G41del_rev | GAGGCCGTCACCATGTCTAAGGGCTCCTCC | SDM |
| DES-S2-A71del_for | CCCCAGCCGGCTCATGGTGACGGC | SDM |
| DES-S2-A71del_rev | GCCGTCACCATGAGCCGGCTGGGG | SDM |
| DES-S2-S81del_for | CTGCGCCGTAGGACATGGTGACGGCC | SDM |
| DES-S2-S81del_rev | GGCCGTCACCATGTCCTACGGCGCAG | SDM |
| DES-S2-F91del_for | CGTCGGCCAGTGACATGGTGACGGCC | SDM |
| DES-S2-F91del_rev | GGCCGTCACCATGTCACTGGCCGACG | SDM |
| DES-S2-F101del_for | TGCGCGTGGTCAGCATGGTGACGGCC | SDM |
| DES-S2-F101del_rev | GGCCGTCACCATGCTGACCACGCGCA | SDM |
| XhoI_Kozak_ATG_DES_S31del_for | TCAGATCTCGAGGCCGTCACCATGTCGCCCGTGTTCCCGCGG | PCR / Cloning |
| XhoI_Kozak_ATG_DES_S51del_for | TCAGATCTCGAGGCCGTCACCATGCGCGTGTACCAGGTGTCGC | PCR / Cloning |
| XhoI_Kozak_ATG_DES_G61_for | TCAGATCTCGAGGCCGTCACCATGGGGGCCGGGGGCCTGGGG | PCR / Cloning |
| BamHI_DES_rev | ACCGGTGGATCCCCGAGCACTTCATGCTGCTGCTGTG | PCR / Cloning |

PCR=Polymerase Chain Reaction; SDM=Site-Directed-Mutagenesis.
